# A foundation model for microbial growth dynamics

**DOI:** 10.64898/2025.12.01.691707

**Authors:** Zachary A. Holmes, Irida Shyti, Alexandra L. Hoffman, Katherine E. Duncker, Helena R. Ma, Zhengqing Zhou, Dongheon Lee, Rohan Maddamsetti, Kyeri Kim, Emrah Şimşek, Grayson S. Hamrick, Hyein Son, César A. Villalobos, Jia Lu, Yuanchi Ha, Ashwini R. Shende, Zhixiang Yao, Sizhe Liu, Daniel M. Shapiro, Kseniia Kholina, Harris Davis, Yasa Baig, Feilun Wu, Shangying Wang, Xiran Wang, Pranam Chatterjee, Michael Lynch, Allison J. Lopatkin, Lawrence David, Emma Chory, Lingchong You

## Abstract

Microbial growth dynamics contain rich information about microbial populations, which support applications from antibiotic testing to microbiome engineering. However, the high dimensionality of growth data and the scarcity of large, task-specific datasets have limited generalizable modeling analysis across systems. Here, we develop a foundation model for microbial growth dynamics. It is a large-scale, self-supervised representation model trained on ∼370,000 experimental and simulated growth curves spanning diverse microbial species, environmental conditions, and community contexts. The model learns lower-dimensional latent embeddings that capture essential dynamical features of raw growth data and enable accurate reconstruction of these data. The concise representations enhance predictive performance in diverse downstream applications. Using these embedding, we achieve few-shot learning for antibiotic classification and concentration prediction, accurate forecasting of simulated and experimental communities, and inference of total abundance from relative-abundance data. By extracting transferable representations from heterogeneous datasets, our model provides a general framework for analyzing and predicting microbial community dynamics from limited measurements.

## Introduction

Microbial growth dynamics are rich in information about how microbial populations behave and interact. For example, growth curves contain information to both identify different strains and predict antibiotic resistance of clinical isolates [1]. A single optical density curve is sufficient to estimate interaction parameters between two strains [2]. Additionally, by predicting microbial consortia dynamics, researchers can design clinically relevant consortia with desired metabolite profiles [3]. Despite their importance, growth dynamics are difficult to generalize across experimental systems and conditions. Variations in strain/species background, assay configuration, and environmental conditions lead to high-dimensional and heterogeneous data that exhibit alternative steady states and abrupt changes to composition [4].

High-dimensional data like these growth dynamics often reside on a lower-dimensional manifold, because many observed features are correlated or redundant [5], [6], [7]. Recent studies show that microbial growth curves can be encoded into lower-dimensional vectors using variational autoencoders, and these embeddings not only allow us to reconstruct the original growth curves but also serve as effective inputs for downstream analyses [8], [9]. Further, microbial growth dynamics are predicted by mapping inputs, such as initial cell densities, ODE parameters, or experimental conditions, to these lower-dimensional vectors [8]. Building on this foundation, we sought to construct a large-scale, generalizable model that learns the universal statistical structure of microbial growth, analogous to how foundation models in other biological domains capture universal relationships within protein sequences [10], genomes [11], or single-cell transcriptomes [12].

Here, we define a foundation model for microbial growth dynamics as a large-scale, self-supervised representation model trained on heterogeneous microbial time-series data to learn generalizable embeddings transferable across tasks and systems. In contrast to models tailored for specific experiments, a foundation model captures shared properties of microbial dynamics, potentially enabling predictions where data are sparse or difficult to standardize. Similar strategies have transformed other fields. For example, the protein foundation model ESM-3 is used to design novel proteins, which simulates evolution [10]. Similarly, the DNA foundation model Evo is trained on genomes and is used to generate functional proteins [11]. Further, the single-cell RNA sequencing foundation model scBERT is trained on RNA sequence databases and can be used to effectively classify different cell types in new samples [12]. We reason that microbial time-series data are likely amenable to such scalable representation learning.

Building a foundation model requires a sufficiently large dataset. Training specific models for tasks in microbial consortia can be inhibitive based on the number of combinations and availability of data [13], [14]. However, there is sufficient data available to train a broadly applicable foundation model. Between high-throughput robotics and collecting previously created growth curves, we curated an extensive dataset on microbial growth [15], [1], [16], [17], [18], [19], [20], [21], [22], [23], [24], [25], [26]. Generating longitudinal studies of microbiota is becoming easier with more access to sequencing technology, and we can collect many of these datasets [4], [27], [28]. Training on broad, diverse datasets leads to better predictive outcomes in downstream tasks [29]. Foundation models have seen a vast array of data, such that they can then be applied to smaller datasets which may be insufficient to train a new model on their own [12], [29].

In this study, we compiled ∼370,000 microbial growth trajectories across multiple species, growth conditions, and community contexts, including both clonal and consortial dynamics. We trained a variational autoencoder on this dataset to learn a compact, low-dimensional latent representation of growth curves. We show that the model accurately reconstructs growth dynamics, generalizes to new datasets, and improves prediction accuracy in data-sparse regimes. Specifically, the model enables few-shot learning for antibiotic classification and concentration prediction, forecasting of microbial consortia dynamics, and inference of total abundance from relative-abundance data. By unifying heterogeneous microbial growth measurements within a shared latent framework, this work establishes a scalable, self-supervised model that serves as a foundation for quantitative prediction and interpretation of microbial population dynamics.

## Results

### Experimental data collection and preprocessing

We collected 306,417 experimental growth curves to build our foundation model. These data consist of:

- Time courses run in a plate reader for a duration of 12 to 40 hours [Fig 1A]
- Continuous cultures run using high-throughput robotics, where the bacteria receive a continuous flow of media throughout the experiment [Fig 1B]
- Longitudinal microbiome studies where the composition is sequenced on a given interval, such as daily over the course of weeks or months [Fig 1C]

**Figure 1.**
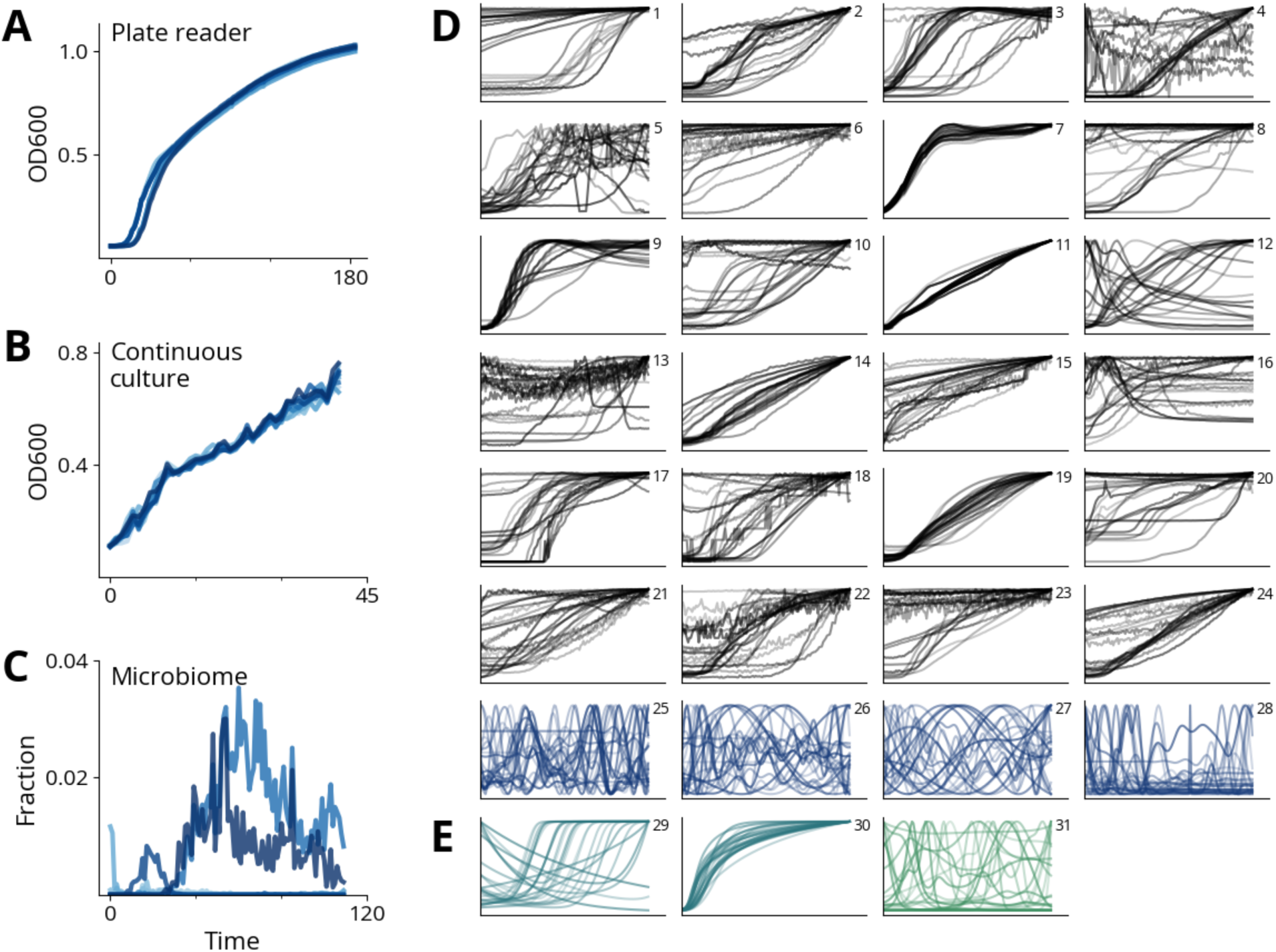
Overview of datasets used to train our model. A) Sample growth curve using OD values from a plate reader.saralaB) Sample growth curve using OD values from a continuous culture reactor.saralaC) Sample growth curves using fractions of strains from a microbiome.saralaD) Experimental growth curves (306,417 curves) were primarily from OD plate readers, automation-enabled continuous cultures, and longitudinal microbiome sequencing. Our data comes from experimental clonal populations (1-24) and experimental consortia (25-28).saralaE) Simulated growth curves (69,674 curves) from clonal populations (29, 30) and microbial consortia (31).

While *E. coli* is the most abundant species in our collected data, the dataset also includes other lab strains, clinical isolates, and natural gut and saliva microbiota, as detailed in the Methods section and supplemental information. We collected both published and unpublished growth data, and we provide a sample of different growth trajectories from the different sources [Fig 1D]. For the plate reader and continuous culture experiments, most of the data contains between 100 to 150 time points, so we selected an input size of 128 time points for our model. Generally, most of the dynamics occur in the first 12 to 20 hours of an experiment. In many cases the experiments were at a steady state by 128 time points. For many of the microbiome datasets, the data were collected on a daily basis. 128 days of growth contain different dynamics than the 12 to 40 hours of growth in the plate readers, so we interpolated the data to bring them to similar scales, despite the diversity of the dynamics. We perform this preprocessing such that our machine-learning model will have consistent input across different experiment types. Details regarding data truncation, interpolation, and preprocessing are included in the Methods section.

### Generation of simulated data

We augmented our experimental data with the simulations of three mathematical models:

- A modified logistic equation
- A generalized Lotka-Voterra (gLV) model with dispersal
- An antibiotic treatment model

The modified logistic equation [Eq 1] stems from the traditional logistic equation and includes additional modifying terms to better replicate the behavior observed in experimental plate reader data, as shown in our previous studies [8], [9], [16]. These modifications account for both empirical growth patterns and background optical density from media, which are not addressed by the standard logistic model. This adaptation enables the simulated data to more closely align with experimental growth curves.

The gLV model with a dispersal term [Eq 2, 3] was adopted from Hu *et al.* [30]. It has been shown to readily generate highly complex dynamics, including chaotic dynamics. We chose the parameters to allow the generation of such complex dynamics.

The antibiotic treatment model [Eq 4-13], based on the work of Ma *et al.*, simulates the dynamics of β-lactam resistant bacteria [16]. It incorporates interactions between resistant and sensitive bacterial populations, as well as the effects of β-lactam antibiotics and β-lactamase inhibitors.

We provide a sample of different growth trajectories from these simulations [Fig 1E].

### Training and evaluating models for universal representation of microbial growth dynamics

Previous work in our lab determined that microbial growth dynamics can be encoded in a lower-dimensional space using machine learning models, such as autoencoders [8]. Here, we developed a foundation model to learn latent representations of microbial growth dynamics. We considered different model architectures, such as custom variational autoencoders, custom transformers, and principal components analysis [Supp Info]. Additionally, we tested the impact of different latent dimensions, ranging from 2 to 32. Based on the results from our analysis, we selected a single model to focus on for the remainder of the paper: the custom variational autoencoder, Autoencoder7X (A7X) with 8 latent dimensions.

A7X encodes 128-time-point curves into an 8-dimensional latent vector, and then decodes the latent vector to reconstruct the original curve [Fig 2A]. Training a variational autoencoder involves computing the mean squared error for the growth curve reconstruction and the Kullback-Leibler divergence loss to enforce a normal latent space distribution. During training, random noise is applied to the 8-dimensional latent vector, with the KL divergence ensuring a Gaussian distribution.

**Figure 2.**
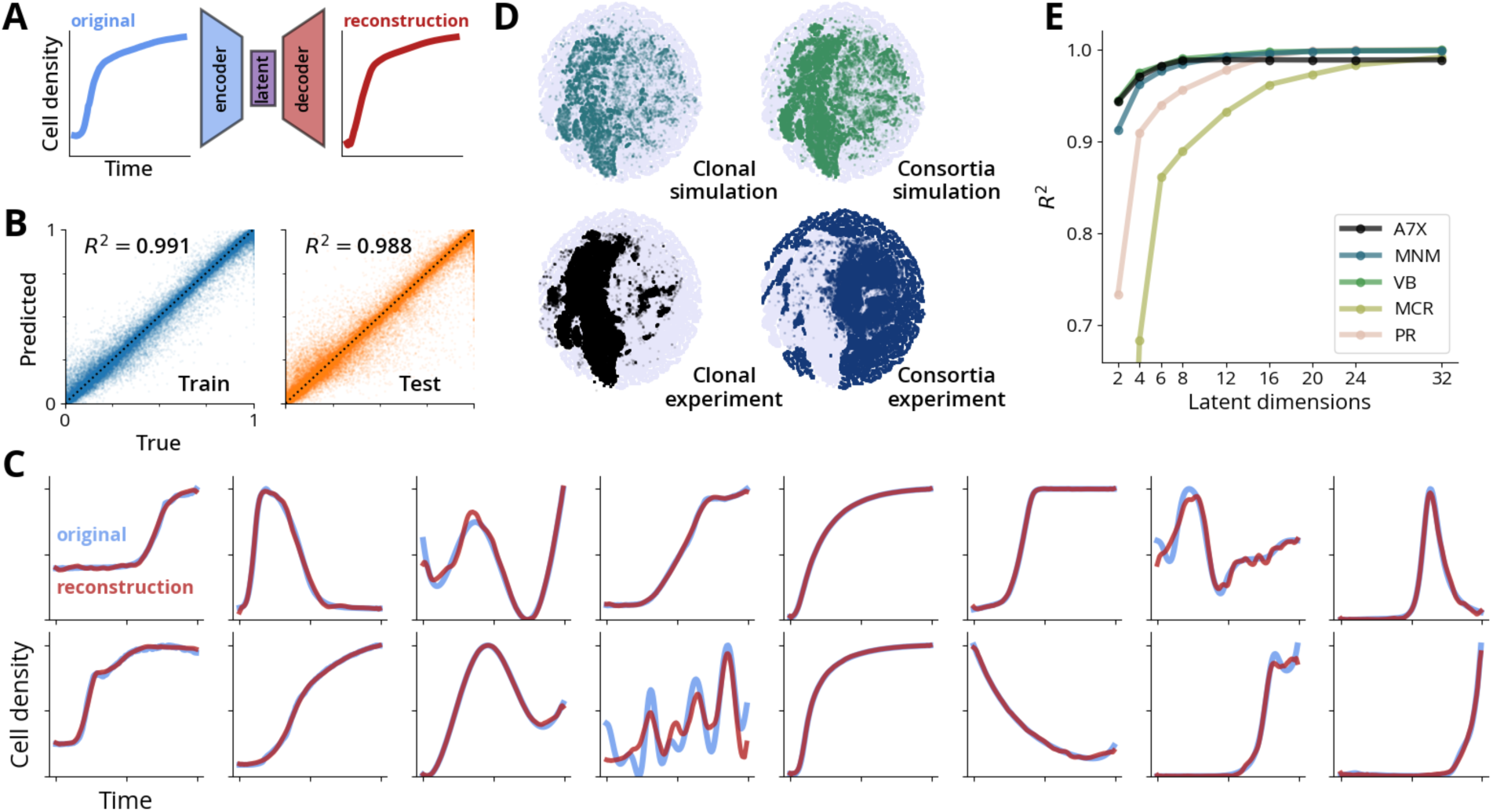
Encoding microbial growth dynamics in a low-dimensional latent space. Our variational autoencoder (Autoencoder7X) compresses a 128-point growth curve into an 8-dimensional latent vector and reconstructs the original curve with high accuracy. A) Schematic of the encoder-decoder architecture used to learn latent representations.saralaB) Reconstruction accuracy for the training and test data. The training dataset is 80% of the total data. The test dataset is 20% of the total data. The model has a coefficient of determination of 0.991 for the training data and 0.988 for the test data. The training and test datasets are 300,872 growth curves x 128 time points and 75,219 growth curves x 128 time points, respectively, which results in 38.5M and 9.6M data points total. For visualization, we randomly selected 100,000 data points from each dataset to display in the scatter plots.saralaC) Examples of original and reconstructed growth curves from our different datasets.saralaD) Visualization of the 8-dimension vectors in 2D by applying t-distributed stochastic neighbor embedding. Different latent values represent different types of growth curves. We visualized where the curves exist in the latent representation for each of the four overarching data types: clonal simulation, consortia simulation, clonal experiment, and consortia experiment.saralaE) Evaluation of different model architectures and latent dimensions. The custom variational autoencoders, Autoencoder7X (A7X), Multi-layer perceptron Network Model (MNM), and Variational autoencoder Bottleneck (VB), have the highest reconstruction accuracy with latent dimensions below 16. Above 16, the principal components analysis reducer (PR) has similar reconstruction accuracy. The transformer architecture, microBERT curve reducer (MCR), does not perform as well, which is to be expected since the transformer is not trained to learn reconstruction, rather it is trained to learn missing data.

We trained our models on a large, diverse dataset: after preprocessing the input data, we used 306,417 experimental and 69,674 simulated growth curves. We decided to supplement the experimental with simulated curves, because including the simulated curves allowed for higher reconstruction of the experimental data for most of the latent dimension sizes [Supp Fig S1]. We trained the model using a training set that consists of 80% of the data for the project (300,872 curves), and 20% was withheld for testing (75,219 curves). Using A7X, the reconstruction of the training data has a coefficient of determination of 0.991, while the testing data is 0.988 [Fig 2B]. The model can accurately reconstruct a broad array of the data [Fig 2C]. We have included more examples in the supplemental material [Supp Fig S2].

We visualize this distribution by encoding all of the curves in our collection and applying t-distributed stochastic neighbor embedding to present the vectors in a 2D display. For each of the four overarching data types - clonal simulation, consortia simulation, clonal experiment, and consortia experiment - we visualized where the curves from those data types exist in the 2D display [Fig 2D].

We find that different model configurations can perform better on different tasks, so the best architecture can depend on a specific application [Fig 2E]. We chose to use a variational autoencoder, because we want accurate reconstructions of growth curves. To select the number of latent dimensions, we used both established statistical methods for estimating intrinsic dimensions, as well as empirical results from our training process. Using the Maximum Likelihood Estimator (MLE) proposed by Levina and Bickel, we estimate the intrinsic dimension of our entire dataset to be 5.6 [7]. As discussed in their paper, estimating intrinsic dimensions is not exact, and the estimated dimension can vary depending on the sample size and parameters used [7]. The estimated intrinsic dimensions vary greatly depending on the complexity of the data, and we see estimates of 1 to 2 for simple growth curves and 8 to 10 for complex or noisy datasets. Therefore, we balance this overall estimate with our empirical observations after testing various dimensions across different models. We selected 8 dimensions based on the accuracy of reconstruction. Latent dimensions below 8 significantly reduced the reconstruction accuracy, while latent dimensions greater than 8 did not add additional benefit to the reconstruction [Supp Table ST5]. The accuracy of the reconstruction is higher when the data come from a less complex dataset, and the accuracy is lower when the data come from a more complex dataset [Supp Fig S3].

### Classifying antibiotic treatment and predicting antibiotic concentration

Few-shot learning is an approach that takes a learned embedding from a foundation model and applies it to a specific problem set [12], [31]. Few-shot learning is especially relevant in microbial biology, because data sparsity is a common problem [13], [14]. Therefore, it is useful to be able to build predictive models using small datasets. We hypothesized that at lower training sizes, the foundation model representation of the curves would outperform the raw growth data. We hypothesized this, because the foundation model reduces the curves to a lower dimension manifold that both reduces noise in the experimental data and emphasizes the key characteristics of the growth curves.

To explore this application, we used a dataset generated by antibiotic resistance work in our lab. This dataset consists of 6,924 growth curves. 672 came from experiments growing *E. coli* TOP10F in different media conditions and initial cell densities [Supp Fig S4]. 5,100 came from experiments growing *E. coli* Keio strains in different antibiotic treatments [Supp Fig S5]. 1,152 came from *E. coli* MG1655 strains grown in different antibiotic treatments [Supp Fig S6]. In our first application, our objective was to classify the type of antibiotic used. In our second application, we predicted the concentration of the antibiotic. For each task, we used a single test dataset, but we adjusted the size of the training dataset to compare the impact of data sparsity on the ability to accurately classify with both the raw curves and the latent vectors.

We trained a classification model to predict which antibiotic was used based on the growth curve data [Fig 3A]. As an input, we used the OD600 curves from the data. We extracted the latent vectors by encoding the curves with Autoencoder7X (A7X). We trained two separate classification models using either the raw curves or the latent vectors as input, and we predicted the antibiotic used. The latent vectors are normalized to a maximum value of one before encoding, so the raw maximum value of the growth curve is appended to the latent vector to give an input vector of 9 dimensions. For this task, we used 4,890 growth curves. These were the OD600 readings for the experimental conditions that had antibiotics added. We used 20%, 978 curves, as the test dataset. For training, we used anywhere from 1% to 80% of the data, 48 to 3,912 curves. We did this with fivefold cross-validation, such that each curve was part of the test dataset in one of the folds. The classification accuracy for the classifier using the latent vectors was higher than the raw data at every training size, and they got closer together with more training data [Fig 3B]. This means that with less data the latent vectors provided more accurate classifications than the raw data.

**Figure 3.**
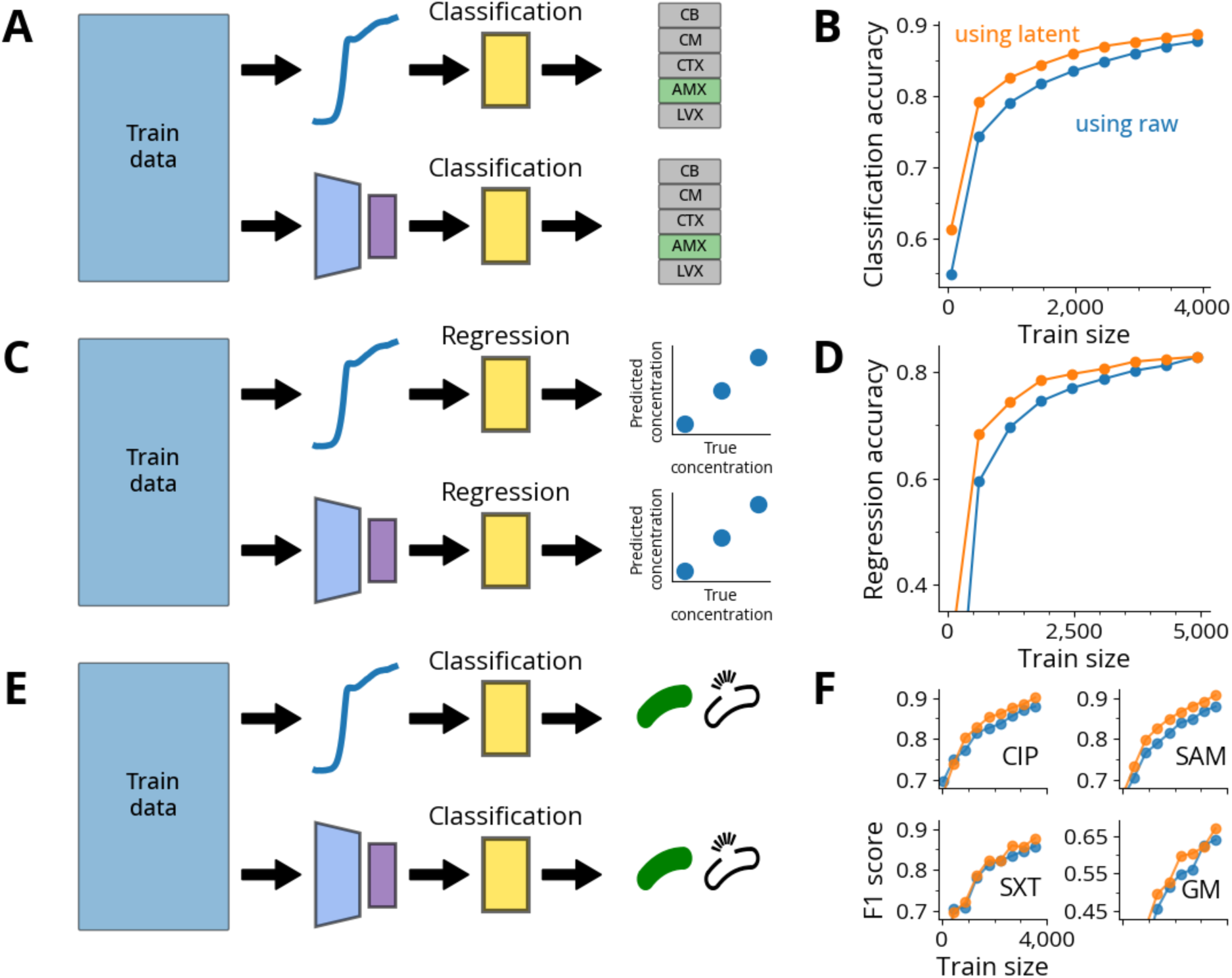
Latent representations enable accurate prediction of antibiotic responses even with a small training sample size. A) Dataset of 6,924 growth curves from *E. coli* strains exposed to different antibiotics and media. In this example, we predicted which type of antibiotic was used to treat the bacteria using only the growth curve as the input.saralaB) Classification of antibiotic type. The latent vectors outperformed the raw curves at every training size, especially under limited data.saralaC) Using the same collection, we also predicted the concentration of antibiotics used to treat the bacteria.saralaD) Prediction of antibiotic concentration by regression. The latent vectors outperformed the raw curves at every training size, especially under limited data.saralaE) Classifying strains. Previous studies in the You lab compared the classification of antibiotic resistance by comparing phenotypes, such as growth curves, against genotypes. Generally, they found using growth curves provided more accurate classifications of strains than the genotypes, even when the growth curves were embedded to a lower dimension [1], [8]. The input data for this task is OD600 readings without any added antibiotics for 277 different clinical isolates, with 4 replicates each in 4 different media conditions, resulting in 4,432 curves. For each antibiotic, we train a classifier on either the raw curves or latent vectors, and we predict whether the cell is resistant to the given antibiotic. Unless otherwise specified, we used the extra-trees regressor and classifier from scikit-learn [42].saralaF1 score for the classifier on each of the antibiotics. With ciprofloxacin (CIP), the latent vectors outperform the raw curves at 7 of the 9 training sizes. With ampicillin–sulbactam (SAM), the latent vectors outperformed the raw curves at every training size. With trimethoprim–sulfamethoxazole (SXT) and gentamicin (GM), the latent vectors outperform the raw curves at 6 of the 9 training sizes. With gentamicin (GM), the latent vectors outperform the raw curves at lower training dataset sizes, but both cases struggle to identify all the GM samples.

We trained a regression model to predict the concentration of antibiotic used in an experiment based on the growth curve data [Fig 3C]. As an input, we used the OD600 curves from the data. We extracted the latent vectors by encoding the curves with A7X. We trained two separate regression models using either the raw curves or the latent vectors as input, and we predicted the antibiotic used. For this task, we used 6,156 growth curves. There were more curves in this task, because we included conditions with no antibiotics added. We used 20%, 1,232 curves, as the test dataset. For training, we used anywhere from 1% to 80% of the data, 61 to 4,924 curves. We did this with fivefold cross-validation, such that each curve was part of the test dataset in one of the folds. The prediction accuracy was higher using the latent vectors than the raw data at every training size, and they got closer together with more training data [Fig 3D]. This was consistent with our hypothesis that less training data will benefit more from the use of the foundation model, but large enough datasets will be sufficient for training the regression model without latent representations. Additionally, we compared the performance of different model architectures and latent sizes on the antibiotic prediction tasks [Supp Fig S8, Supp Table ST6, Supp Table ST7].

### Classifying antibiotic resistance from cell density curves

Bacterial growth curves contain sufficient information to predict phenotypes such as antibiotic resistance to ampicillin–sulbactam (SAM), ciprofloxacin (CIP), trimethoprim–sulfamethoxazole (SXT), and gentamicin (GM), as shown in previous work from our lab [1]. Predicting these phenotypes from growth data is meaningful, because 1) genotypes do not have a direct linkage to phenotypes, so WGS is insufficient for making these predictions and 2) we can use a simple OD600 growth curve which does not require testing different antibiotics individually [1]. In previous work from our lab, Baig *et al*. trained a variational autoencoder end-to-end on a subset of these curves, and by embedding the derivatives of the curves they were able to classify the resistance more accurately than with the raw data [8]. Following their work, we performed a similar task of classifying the resistance of bacteria based solely on their OD600 growth curves. Instead of taking the derivative of the curve, we used the blank-corrected data as the input.

There were 4,432 growth curves in this dataset [Supp Fig S7]. The dataset consisted of 277 clinical isolates with 4 growth conditions, and each experiment had 4 replicates. Of the 277 strains, 42 are resistant to none of the antibiotics, 47 are resistant to 1, 114 are resistant to 2, 45 are resistant to 3, and 29 are resistant to all 4. We used a training size of 80% of the data randomly separated, so the training dataset was 3,324 curves and the test dataset was 1,108 curves. We did this with fivefold cross-validation, such that each curve was part of the test dataset in one of the folds. We tested 2 different input configurations to predict the resistance: raw 128-time-point curves and 8-dimensional latent vector from Autoencoder7X (A7X) foundation model encoding [Fig 3E]. The latent vectors are normalized to a maximum value of one before encoding, so the raw maximum value of the growth curve is appended to the latent vector to give an input vector of 9 dimensions. No information about the strains or growth conditions were included in the model.

For each of four types of antibiotics, we ran our training pipeline to classify whether a given strain was resistant or sensitive to the antibiotic. The results are reported as the F1 score, which is a balance of precision, the ability to avoid false negatives, and recall, the ability to identify all of the positives. We found that for two of the antibiotics, SAM and CIP, the latent vectors outperformed the raw data [Fig 3F]. For the other two antibiotics, SXT and GM, the latent vectors and raw data are comparable [Fig 3F]. The foundation model provides the benefit of having seen many more growth curves than what is available in this study, and this is why its latent vectors are able to meaningfully represent the growth curves in this sparse dataset. Additionally, we compared the performance of different model architectures and latent sizes on the antibiotic prediction tasks [Supp Fig S8, Supp Table ST8, Supp Table ST9, Supp Table ST10, Supp Table ST11].

### Forecasting simulated microbial community dynamics

Microbial consortia are found everywhere in nature: from human saliva, skin, and guts to soil or the ocean [4], [27], [28]. Researchers often want to predict the behavior of microbial consortia to understand the impact on things like metabolite production in the gut [3]. While understanding how an entire microbial consortia operates can be important, in many cases we are only interested in a few key members of the consortia. For example, when examining complex microbial consortia, we typically focus on a few key strains, such as the infectious gut pathogen *Clostridioides difficile* and the inhibitory strains *Clostridium scindens* and *Clostridium hiranonis* [32]. In this case, we can modify our dataset to focus on the key members of a consortium and omit the background strains and consider their influence as background noise. Using our foundation model, we can represent this community in a lower dimensional vector and predict its dynamics and behavior into the future.

For this application, we are using a dataset which consists of 150 different consortia [Supp Fig S9]. These consortia were generated using two different variations of the generalized Lotka-Volterra model. The first is the dispersal variation [Eq 2, 3] used by Hu *et al.* [30]. The second is the bounded version [Eq 14] published from our lab [33]. These models result in both stable and chaotic behavior. Each of the consortia was originally run with 20 to 100 members in the community. From these original simulations, 2 to 10 focal members were selected for the analysis. Each consortia was simulated 10,000 times, unless stated otherwise, and we split the data randomly into 8,000 curves for the full training set and 2,000 curves for the test set. In each simulation, the initial quantities of the focal populations were randomized. For some of the consortia, the background communities had the same initial quantities in each simulation, and in others they were randomized. From the 8,000 curves in the training set, we randomly selected smaller training sizes of 20, 100, 200, 1,000, and 4,000 curves. On each of our 6 training datasets, we performed fivefold cross-validation to train 5 separate models, and we report the mean values from the results of these models. All final metrics are measured on the original test set with 2,000 curves.

To demonstrate the utility of our model, we selected two of these consortia to analyze. The first community is our simple community with 3.0 estimated intrinsic dimensions. This consortium was generated using 20 background strains, and we randomly selected 5 members as the focal community. The second community is our complex community with 6.0 estimated intrinsic dimensions. This consortium was generated using 100 background strains, and we randomly selected 8 members as the focal community.

We hypothesize that these curves have sufficient content to predict future behavior of the consortium. We designed a pipeline to train a regression model to use 128 time points to predict the 128 time points [Fig 4A]. For each of the input configurations, the curves were stacked such that the input to the regression model with *M* communities x 128 or 8, respectively. We tested 2 different input configurations.

1) Raw 128-time-point curves
2) 8-dimensional latent vector from Autoencoder7X foundation model encoding plus maximum value of raw curve for a total size of 9 dimensions

**Figure 4.**
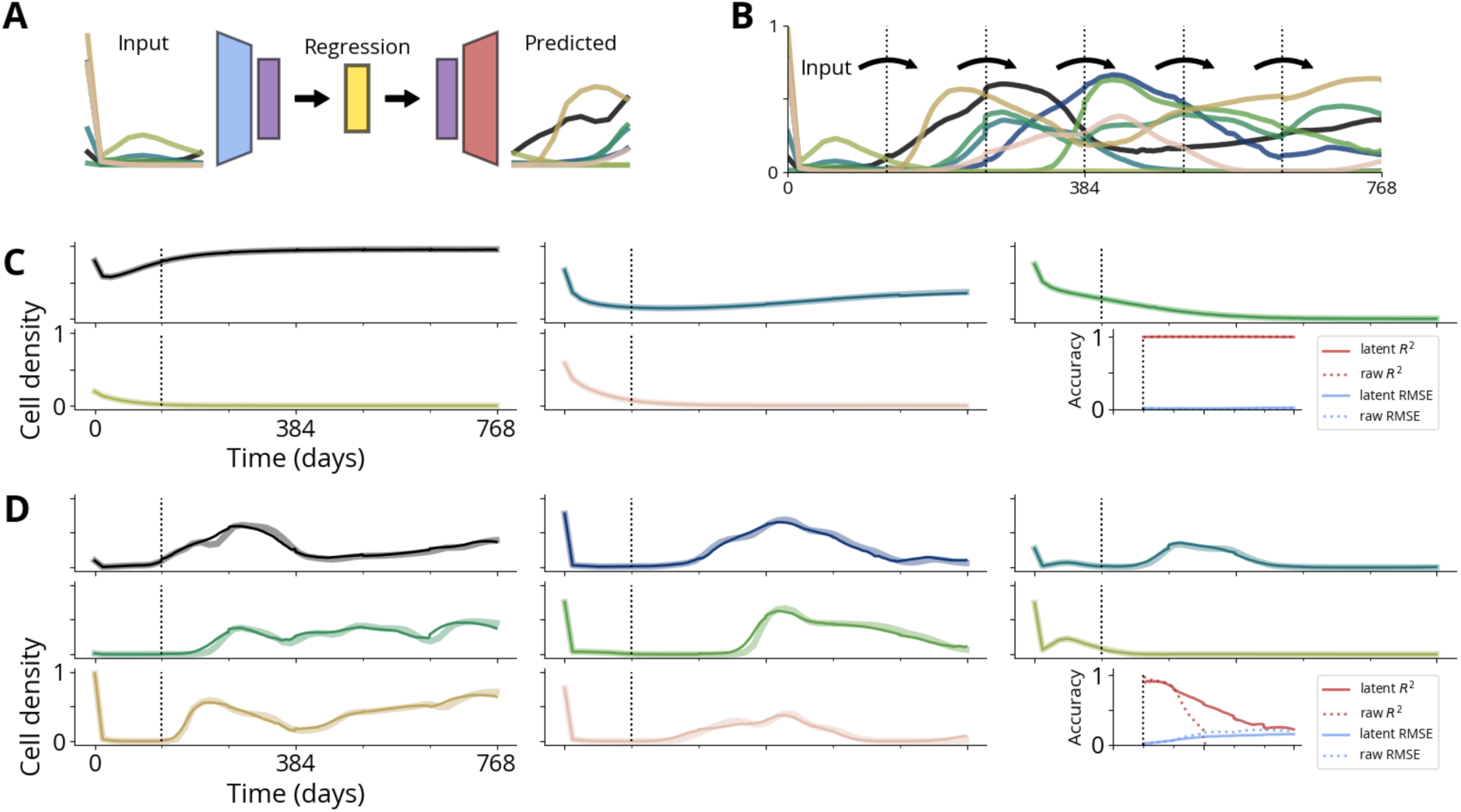
Forecasting simulated microbial community dynamics using latent representations. A) Workflow for training a regression model for forecasting. We defined 150 consortia and simulated 10,000 curves for each consortium. The consortia each contained 20 to 100 members. For each consortium, we selected *M* (= 2 to 10) members for our analysis (see Methods for details). From the simulated data, we use 128x*M* time points to predict the next 128x*M* time points. We tested two different conditions for this task: 1) using the raw data to predict the raw data and 2) using the latent representation of the input to predict the latent representation of the output. For the latter, our pipeline consists of training the regression model by providing the input curves, encoding to the latent representation, training the regression model to predict the latent representation, which is then decoded into the future 128 time points. We split the 10,000 curves into 8,000 for the training dataset and 2,000 for the test dataset.saralaB) Iterative forecasting using the trained regression model. Using a single input of the first 128 time points from the simulation, we predict the remaining 640 time points. The regression model is trained to take 128 time points and predict the next 128 time points. Therefore, we use points 1 to 128 to predict 129 to 256. Then, we use the newly predicted 129 to 256 to predict 257 to 384. We perform this iteratively until we have predicted the entire time course.saralaC) Forecasts for a simple consortium (estimated intrinsic dimension ≈ 3) using latent representations. The displayed results are from using 10% of the training data, or 800 simulations, to train a regression model. This community was simulated using 20 total members, and we selected 5 as target members. The wider, faint lines are the ground truth and the skinnier, solid lines are the forecasted behavior. Each color is a different member of the community. The displayed example is a random sample from the 2,000 test simulations. Regression accuracy of R^2^ (red) and RMSE (blue) for both using raw (dotted) and latent (solid). These metrics are calculated by taking the mean values from fivefold cross-validation from training regression models on the 800 curves. For this simple community, both using raw and latent perform very well, and there is no difference in the accuracy between the two.saralaD) Forecasts for a complex community (estimated intrinsic dimension ≈ 6). These are 8 members from a total community of 100 members. The displayed results are from using 10% of the training data, or 800 simulations, to train a regression model. The displayed example is a random sample from these 2,000. Regression accuracy of R^2^ (red) and RMSE (blue) for both using raw (dotted) and latent (solid). For the complex community, we see that using latent outperforms using raw significantly in the latter half of the forecast. These metrics are calculated by taking the mean values from fivefold cross-validation from training regression models on the 800 curves.

Inspired by the success of GraphCast and Lam *et al.* in forecasting weather [34], we use our regression models to forecast microbial consortia behavior into the future. We use an initial seed of 128 time points, and then we predict the next 128 time points by using the latent vector as the target value. We then append the predicted results to the initial seed, and slide the window to use the updated prediction as the input for the next segment [Fig 4B].

For the simple consortium, we are able to accurately forecast into the future [Fig 4C]. To demonstrate the utility of the foundation model at lower training data sizes, we used the training dataset with 800 curves. To generate the accuracy metrics, we used each of the 5 models from the fivefold cross-validation and used them to forecast the 2,000 curves from the test dataset. The reported metrics are the mean values of these 5 forecasts. The accuracy of the prediction is high with a coefficient of determination above 0.995 and RMSE below 0.022 for both raw curves and latent vectors. For the complex consortium, we forecast all the members into the future [Fig 4D]. The forecast accuracy decreases over time, but the latent vectors provide better accuracy further into the future compared to the raw curves as inputs. We have included more examples in the supplemental material [Supp Fig S10].

### Forecasting experimental microbial community dynamics

While predicting simulated consortia is a useful exercise, being able to predict real-world microbial consortia is a much more meaningful task. For this task, we use the microbiome experiments completed by Fujita *et al.* on their natural microbial communities [4]. This dataset is useful in our analysis, because they have 48 unique experiments. The experiments are completed using 2 different source microbiomes in 3 different media types with 8 replicates each [Fig 5A].

**Figure 5:**
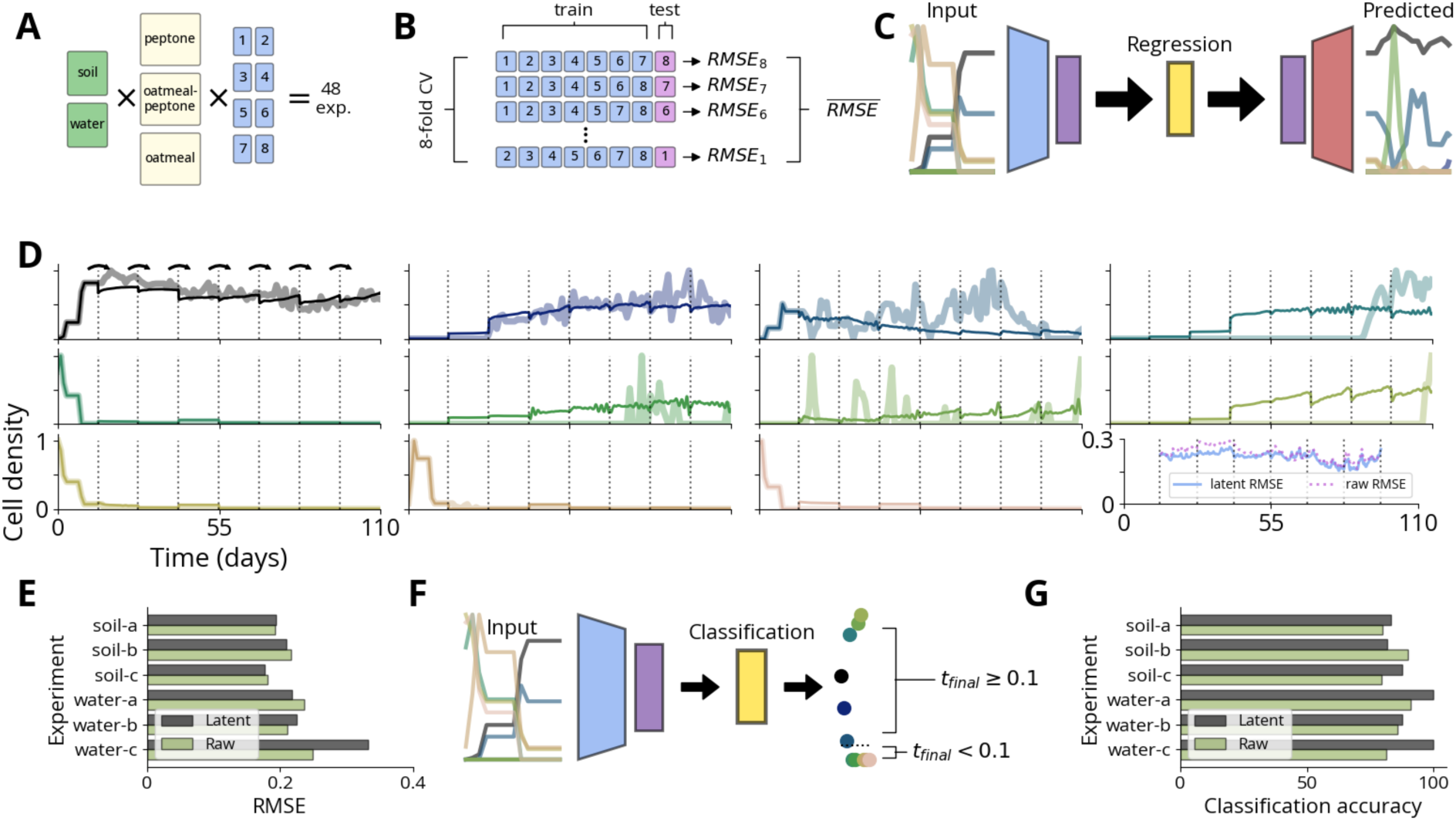
Forecasting and abundance prediction in experimental microbial consortia. A. Experimental data, from Fujita *et al.* 48 consortia derived from two environmental sources were cultured in 3 different media types, each with 8 replicates and measured daily for 110 days [4]. Each of the source/media combinations had 28 to 144 members, however not every member was present across all 8 replicates. For each source/media combination, we selected the members (2 to 18) that were present across all 8 replicates as our focal members.saralaB. Eightfold cross-validation treats each replicate as an independent test set to quantify predictive performance. We report our results by averaging the output of the model, such as taking the mean RMSE for predicting the consortia behavior.saralaC. Regression pipeline for forecasting. The model uses 128x*M* time points as input to predict the next 128x*M* time points, where *M* is the number of selected members. We consider 2 different methods: 1) using raw curves to predict raw curves and 2) using latent representations to predict latent representations. For each iteration of the cross-validation, we are using 7 replicates for the training. The original data consists of 110 daily readings. We interpolate this data to a total length of 768 time points, which results in about 14 days per 128 time points. During training, we take every slice from these training curves: we use time points 1 to 128 to predict 129 to 256, 2 to 129 to predict 130 to 257, 3 to 130 to predict 131 to 258, etc. This results in a training set of 512 samples per replicate, or 3,584 samples total.saralaD. Example forecasts for one consortium. Each panel is one of the 11 members from the consortium. RMSE for these data (final panel) is much higher than the simulated data from the previous figure. For this consortium, latent forecasts (solid) are slightly better than those based on the raw curves (dotted).saralaE. Across six source/medium combinations, latent-based forecasting yields lower mean RMSE than raw-based forecasting in half of the conditions. Overall, the two approaches perform similarly.saralaF. Predicting end-point dominance using early-time-window data. We take our initial 128 time points, which equates to 14 days, and predict whether the final abundance on day 110 will exceed 10%. Similar to the forecasting, we use eightfold cross-validation and report the mean accuracy from the results.saralaG. Latent vector representation of the input data leads to higher classification accuracy in 5 of the 6 experimental conditions.

One challenge with collecting microbiome data is the quality of the data. Species and members in a consortia that are lower in abundance are harder to measure, so we only use members that are above a certain abundance for this task. For each of the 6 experimental conditions, there were anywhere from 6 to 77 strains present in the experiment. For each condition, we selected 2 to 18 members that were present in all 8 of the replicates [Supp Fig S11]. We treat these as our focal communities within the larger consortia. We applied eightfold cross-validation and used each of the replicates as the test set for each round of validation [Fig 5B]. This allows us to measure the average performance of prediction on the community instead of relying on a single sample.

Similar to the simulated consortia, we designed a pipeline to train a regression model to predict the next segment of points based on the previous 128 time points [Fig 5C]. For each of the configurations, the curves were stacked such that the input to the regression model with *M* communities x 128 or 9, respectively. We tested 2 different configurations.

1) Input of latent vector plus maximum raw value to predict latent vector plus maximum raw value of the next 128 time points
2) Input of raw curve to predict raw curve of the next 128 time points

As with the simulated consortia, we forecasted the behavior of the focal communities [Fig 5D]. We use an initial seed of 128 time points, and then we predict the next 128 time points. Forecasting the experimental consortia is more difficult than the simulated consortia, and we find that despite the latent to latent prediction outperforming the raw to raw prediction, the minimum RMSE value across all the experiments is 0.177 [Fig 5E]. We have all examples in the supplemental material [Supp Fig S12].

We expand this application to a task that is useful in analyzing microbial consortia: predicting abundance in the future. For all the communities, we used the first 128 time points to predict which species would have an abundance above 10% at the final time point. In other words, we use the first 14 days worth of interpolated data to predict the species abundance 96 days later [Fig 5F]. We find that the latent input outperforms the raw input in 5 of the 6 experimental conditions [Fig 5G].

### Predicting absolute abundances from relative abundances

In microbiome research, we are generally interested in understanding the composition of microbial consortia [35]. A common way to derive composition is to use amplicon sequencing, such as 16S rRNA, which provides relative abundance of different members of a consortium [36]. However, these measurements lack information about total microbial abundance, which is critical for understanding ecological dynamics, host-microbe interactions, and community stability [35], [37], [38]. We hypothesize that temporal patterns in relative abundance data contain sufficient information to infer total abundance dynamics. Furthermore, we propose that foundation models trained on diverse microbial growth dynamics could extract features from these temporal patterns, and these features enable more accurate predictions than using raw relative abundance data alone.

To test our hypothesis, we generated two distinct synthetic datasets, each containing 10,000 simulated microbial communities with 20 species tracked over 128 time points. The first dataset modeled community dynamics using a gLV variant that incorporated logistic growth, saturating positive interactions, and density dependent negative interactions [33]. The second dataset was based on a classical generalized Lotka-Voltera model with strong intraspecific competition and a weak uniform dispersal term, designed to capture more complex and potentially chaotic dynamics. Both models used randomly sampled interaction parameters to create diverse community behaviors, with initial conditions drawn from a Dirichlet distribution to ensure varied compositional starting points at low total biomass. From each absolute abundance trajectory generated by these models, we computed relative abundances by normalizing by the total community abundance at each time point, creating paired datasets where the task was to predict total abundance from relative abundance profiles.

For each sample, we used the complete trajectory of relative abundances across all species (20 species × 128 time points) as input to predict the corresponding total abundance at each time point. We compared two input representations:

1) Raw 128-time-point curves of relative abundance data for each of the 20 members [Fig 6B]
2) 8-dimensional latent vector from Autoencoder7X foundation model encoding plus maximum value of raw curve for a total size of 9 dimensions for each of the 20 members [Fig 6A]

**Figure 6.**
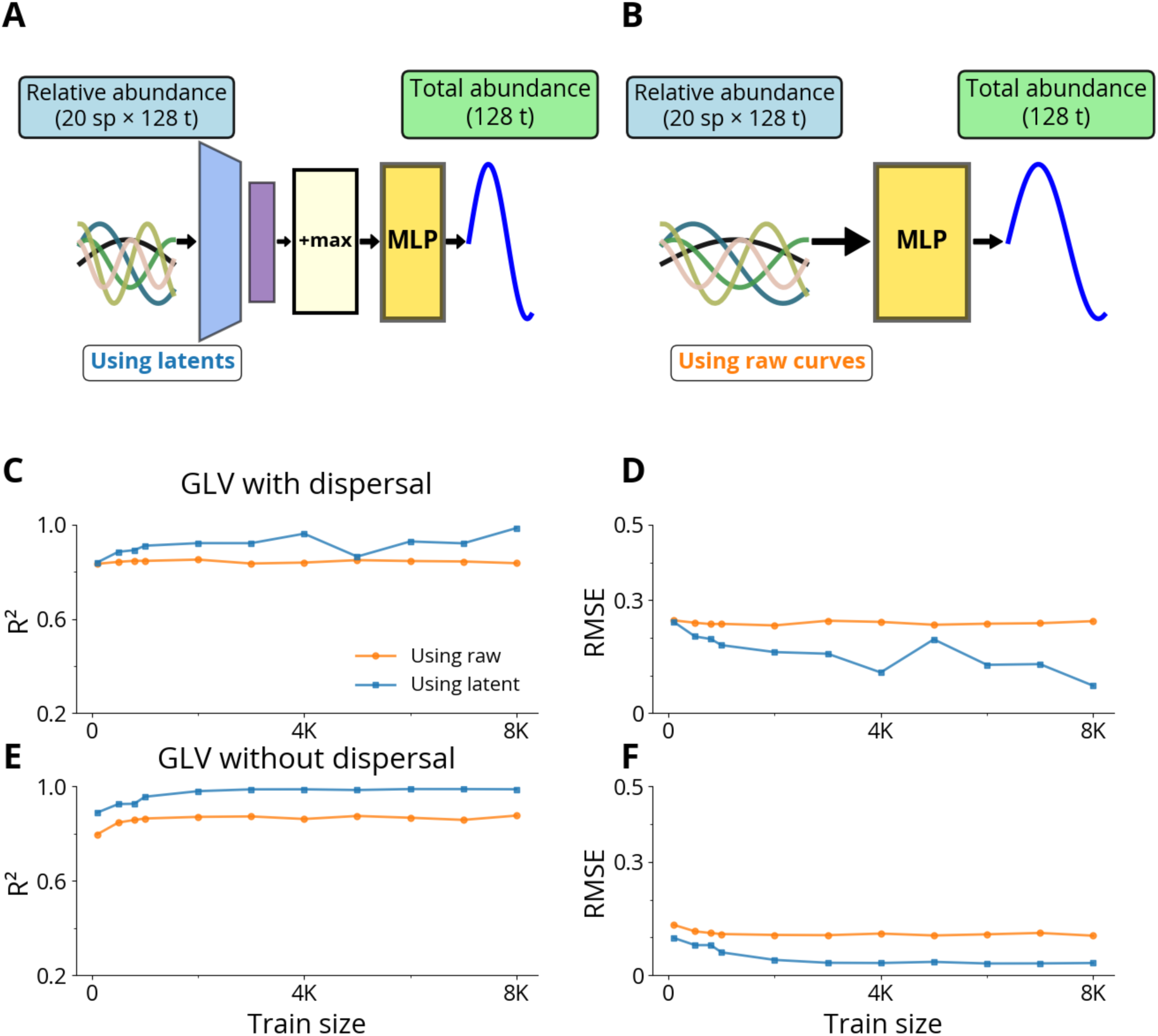
Predicting total abundances from relative abundances in simulated communities. A. Relative abundance trajectories (20 species × 128 time points) are encoded by the foundation model (Autoencoder7X) into an 8-dimensional latent space. This compressed representation is concatenated with the maximum relative abundance value to create a 9-dimensional feature vector, which is fed into an MLP regressor to predict total abundance dynamics (128 time points). Example curves illustrate diverse temporal patterns in species abundances and the predicted output trajectory.saralaB. The same relative abundance input data is directly fed into an MLP regressor without dimensionality reduction or feature extraction (2,560 total input dimensions: 20 species × 128 time points). The model outputs predicted total abundance trajectories using the same prediction framework as the latent pathway.saralaC. R² for both approaches evaluated across increasing training set sizes on complex community dynamics generated from a generalized Lotka-Volterra framework. Latents consistently outperform raw data across all training sizes.saralaD. RMSE values for both prediction approaches show that using the latents achieves lower prediction error than using the raw data across all training set sizes, with the performance gap particularly pronounced at smaller training set sizes.saralaE. For stable GLV dynamics, both raw relative abundance data and foundation model latent representations achieved nearly identical performance, suggesting that the relatively simple, predictable dynamics could be captured effectively by either representation.saralaF. RMSE values for the stable dynamics regime show comparable prediction errors between the latent and raw pathways across all training set sizes.

We implemented multi-layer perceptron (MLP) regression models to predict total abundance from the input features. Both models directly predicted 128-dimensional output vectors corresponding to total abundance at each time point, using mean squared error as the loss function. Critically, for the latent representation approach, we bypassed the foundation model’s decoder: the MLP predicted total abundance directly from the compressed latent features without reconstructing back to the original data space. This design allowed us to test whether the latent representations captured dynamics-relevant information sufficient for prediction, independent of reconstruction accuracy. We trained both models using fivefold cross-validation across varying training set sizes (1% to 80% of data), implementing early stopping based on validation loss to prevent overfitting [Fig 6A, Fig 6B].

The results demonstrate that foundation model latent representations consistently outperform raw relative abundance data across both dynamical regimes. For the GLV without dispersal dynamics, the latent representation achieved superior performance across all training set sizes [Fig 6E]. The GLV with dispersal dynamics showed the same patterns [Fig 6C]. The foundation model’s compressed latent space (9 dimensions) capture more generalizable dynamical features that are transferred effectively to the prediction task compared to the high-dimensional raw input space. We observe similar trends for RMSE: latent features consistently outperform raw data for both GLV with dispersal [Fig 6D] and GLV without dispersal [Fig 6F]. This finding supports our hypothesis that foundation models pre-trained on diverse microbial dynamics extract robust, transferable features that enable better generalization than raw temporal patterns, with consistent advantages across both simple and complex dynamical regimes.

## Discussion

Microbial growth trajectories are among the most fundamental and widely measured descriptors of population behavior. Yet, despite decades of data, the lack of a unified analytical framework has limited our ability to compare, interpret, or predict growth dynamics across species, environments, and community contexts. Here, we introduce a foundation model for microbial growth dynamics. This model is a large-scale, self-supervised representation model that learns the shared statistical structure of microbial time series from heterogeneous datasets. The resulting latent representations capture key dynamical features, which enable accurate reconstruction, transfer across experimental systems, and improve performance in downstream predictive tasks.

Our model builds on the recognition that microbial dynamics, while high-dimensional in their raw form, lie on a low-dimensional manifold. Previous studies have established this empirically within the context of specific systems and conditions [8], [9]. Our current study extends that insight to a far broader scale, integrating hundreds of thousands of experimental and simulated trajectories to reveal the universal structure underlying microbial population dynamics. This generalization represents a conceptual advance: rather than fitting a single mechanistic model to each system, we train a single data-driven model that captures the statistical essence of microbial growth across systems and conditions.

Similar to protein or transcriptomic foundation models, which learn universal embeddings from large-scale biological data, our model distills microbial time-series behavior into compact latent variables that can be reused for diverse applications, including classification, forecasting, and inference, from limited or noisy measurements.

The success of this framework reflects both biological and statistical regularities. Biologically, microbial populations often exhibit stereotyped phases of lag, exponential growth, and saturation, which constrain the effective dimensionality of the system. Statistically, these constraints allow a deep encoder-decoder architecture to learn transferable representations that summarize population-level behavior. The emergent low-dimensional structure suggests that microbial growth, despite the complexity of underlying physiology, can be effectively characterized by a small set of latent variables.

The model, along with the training, can be revised and expanded in multiple ways. More data can be included in the training process. This would involve sourcing from across all publications to include as much data as possible, and continuing to collect data from our colleagues and collaborators. Different model architectures could allow for more flexibility, or may perform better in different tasks. The requirement for 128 time points requires truncation or interpolation, which is one potential limitation in this approach. Designing a model to take an input of any size would allow for more flexible applications. Additionally, actual time between samples is not a focus in this study. Future versions of this model should include a time component that allows the model to interpret specific time points from experiments. This could overcome some of the limitations we saw with trying to represent experiments with 5-minute gaps between data in the same model as daily readings from the gut. Finally, future versions could incorporate categorical data, such as strain information, media type, and experimental conditions, so the model could learn how these known factors influence growth.

Beyond microbiology, our work highlights the broader potential of large-scale, self-supervised learning to extract interpretable and generalizable representations of biological dynamics. The ability to encode population behavior into transferable latent features provides a bridge between data-driven discovery and mechanistic modeling, complementing rather than replacing explicit models of growth, metabolism, and ecological interaction. By establishing a unifying latent framework for microbial time series, our study offers a conceptual and practical foundation for predictive microbiology, where data collected in one context can inform inference, design, and control in another.

## Methods

### Plate reader experiments

Plate readers are used to measure optical density and fluorescence over time. Optical density corresponds with the biomass of the cells in the plate wells. Fluorescence is a combination of fluorescence output per cell and the number of fluorescent cells. The general protocol is as follows. Cells were inoculated the night before the plate reader experiment began and grew to stationary phase. The next morning, the cells were diluted and added to media and chemical combinations according to the specific experimental protocol. These mixtures were added to plates, which were placed in the plate reader. The plate reader maintained an experimental temperature, such as 37°C. Every 5 to 10 minutes, as specified by the experiment, the plates were shaken and then measurements were taken. This resulted in growth curves as measured by optical density and fluorescence. The exact experimental conditions, including media, treatments, temperatures, inoculum densities, etc., varied depending on the data source, and further details can be found in the Supplemental Information.

### Continuous culture experiments

Continuous culture experiments were completed using liquid handling robots. There were two methods used for these experiments: turbidostats or basic kinetic experiments. The high-throughput turbidostats were bacterial wells continuously fed with media. The real-time feedback allowed the robot to adjust the flow to the turbidostat based on the measurements of the growth of the cells, which enabled calculating the growth rate of the cells. A detailed protocol on the implementation of this system can be found in the publication from Chory *et al.* [39].

### Microbiome experiments

Microbiome experiments are conducted by sampling microbiome communities and running sequencing to quantify the abundance of each of the members in the community. We used four different collections of microbiome data in our study.

1) Microbiomes were collected from natural samples and experimented with different media conditions [4]. We used the data directly from the published article.
2) Gut and saliva samples [27].
3) Our lab has previously developed a collection of bar-coded Keio strains, and we used experimental data from communities of these strains [33], [40].

### Experimental data preprocessing

All original growth curves are available in the supplemental data. While many of the growth curves were previously published, this study also includes new growth curves generated under various configurations, as described in the supplemental material [Supp Table ST12].

For plate reader and continuous culture datasets with more than 128 time points, we truncated the data after 128. For experiments with less than 128 time points, we interpolated to fill in the missing data. We interpolated time courses from the plate reader using linear interpolation.

Microbiome datasets presented additional challenges due to their longer timescales. To address this, we segmented the microbiome data into 16-day chunks. Missing data points were interpolated using a Gaussian Process instead of linear interpolation, as Gaussian Processes produced smoother, organic shapes that better captured the expected trends in these datasets.

The data were combined into a single dataset of 376,091 curves. The data were split into a training set of 80% of the curves and a test set of 20% of the curves. We split the data by iterating through the dataset and randomly splitting every 1,000 curves into 800 for training and 200 for test, so that each dataset was properly represented in each group.

### Simulation using a modified logistic equation

Our lab developed a modified logistic equation to address the limitations of the traditional logistic equation, which does not adequately replicate growth curves from plate readers [8], [16]. Plate readers are subject to background optical density from the media as well as the secondary growth stages. To better fit the logistic function to experimental data, we introduced two modification constants: *k* and *θ*.

In the following equation, the variables are as follows:

- *n* is the cell density
- *t_lag_* is the lag time
- *⍺* is the cell growth rate
- *n_max_* is the carrying capacity
- *k* is a modification constant
- *θ* is a modification constant

Simulation parameters are provided in the supplemental data [Supp Table ST1].

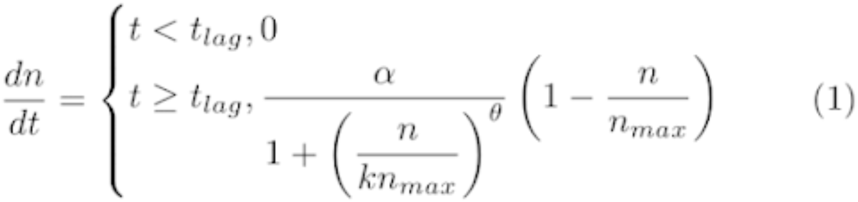

### Simulating community dynamics

To generate dynamics with varying complexity (including chaotic dynamics), we used a generalized Lotka-Volterra model with dispersal, as published by Hu *et al.* [30]. The model was chosen for its ability to capture complex, dynamic interactions within microbial communities. Specifically, we focus on chaotic interactions, because they provide more examples of cells with changing cell density, as opposed to the examples with stable steady states.

In the following equations, the variables are as follows:

- *n* is the cell density
- *A* is the interaction matrix
- *D* is the dispersal rate
- *a_i,j_* is the interaction of species *i* with species *j*
- *M* is the number of members in the population

Simulation parameters are provided in the supplemental data [Supp Table ST2].

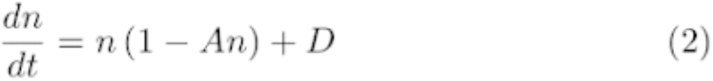

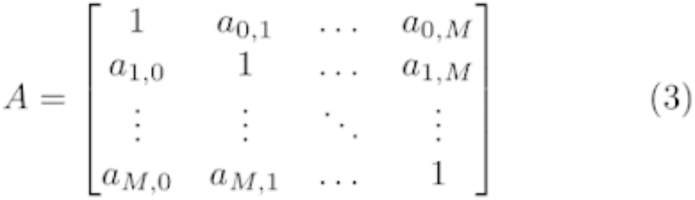

### Simulating growth dynamics in response to *β-lactam treatment*

To simulate antibiotic resistance, we used the β-lactam resistance model described by Ma *et al.* [16]. This model employs ordinary differential equations to capture the dynamics between resistant and sensitive cells and their response to β-lactam antibiotics and β-lactamase inhibitors.

In the following equations, the variables are as follows:

- *n_s_* sensitive cell density
- *n_r_* resistant cell density
- *s* nutrient concentration
- *a* antibiotic concentration
- *b* β-lactamase concentration
- *g* growth rate
- *l* lysis rate of cells
- *γ* lysis rate by antibiotic
- *⍺* cost of β-lactamase production
- *β* benefit of β-lactamase production
- *ξ* nutrient recycling
- *φ* antibiotic degradation by living cells
- *i* β-lactamase inhibitor concentration
- *d_b_* effect of β-lactamase inhibitor on β-lactamase
- *c* private benefit
- ***κ****_b_* antibiotic degradation by β-lactamase
- *d_a_* basal antibiotic degradation
- *h_a_* Hill coefficient for antibiotic
- *h_i_* Hill coefficient for inhibitor
- *ι* effect of β-lactamase inhibitor on β-lactamase

Simulation parameters are provided in the supplemental data [Supp Table ST3].

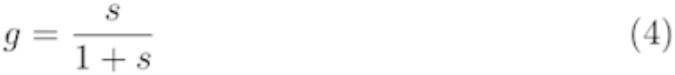

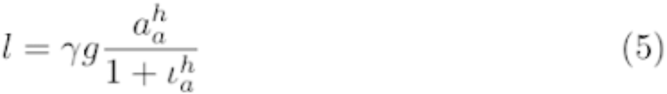

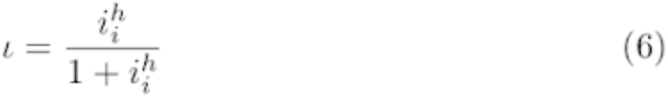

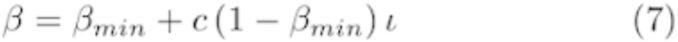

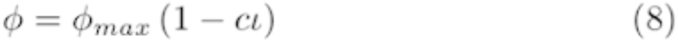

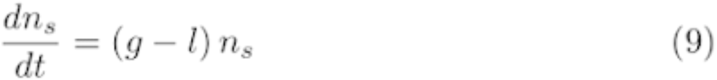

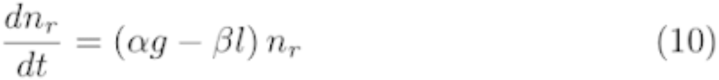

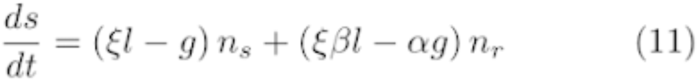

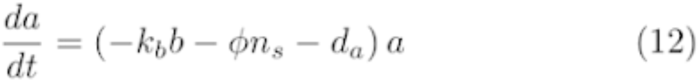

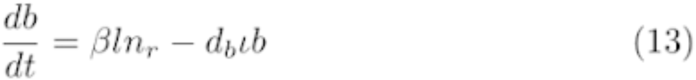

### Simulation using a bounded generalized Lotka-Volterra

To generate consortia bounded between values of 0 and 1, we used a generalized Lotka-Volterra model, as published by Wu *et al.* [33]. The model was chosen for its ability to generate meaningful consortia dynamics.

In the following equations, the variables are as follows:

- *n* is the cell density
- *μ* is the growth rate
- *σ* is the environmental stress
- *γ*^+^ is the positive interaction matrix
- *γ*^−^ is the negative interaction matrix

Simulation parameters are provided in the supplemental data [Supp Table ST4].

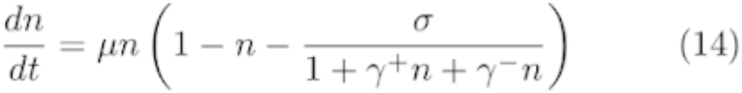

### Foundation model formulation

Our custom variational autoencoder, Autoencoder7X (A7X), consists of 7 encoding layers and 8 decoding layers. For the encoder, the first 6 layers are convolution layers which convolutes the 128-time-point curve to a 128 x 8 vector. The vector is then fed into the final layer of the encoder, which consists of using a linear layer to reduce the vector to the latent dimension size 8. During training, this final layer of the encoder consists of both a mu linear layer that calculates the mean values of the latent vector, and a logvar linear layer that applies random noise to the mean. Therefore, each input corresponds to a continuous latent range, and not a single value. This noise is regularized to a Gaussian distribution using the Kullback-Leibler divergence loss. During evaluation, only the mu layer is used to compute the latent vector. The decoder architecture reverses the encoder. It first takes the 8-dimensional latent vector and passes it through a linear layer. Then, it performs 7 transposed convolutions to output a 128-dimensional vector. The decoder is made up of 8 layers. The first layer is a linear layer which receives the latent vector as the input. The remaining layers are transposed convolution layers. The last layer outputs the reconstructed growth curve. The losses were calculated using both mean squared error and Kullback–Leibler divergence. For each model architecture, we used Optuna to identify optimal hyperparameters [41]. Optuna is a framework developed to optimize hyperparameters in machine learning tasks.

### Antibiotic data preprocessing

For each of the three antibiotic tasks, 1) classifying treatment, 2) predicting concentration, and 3) classifying resistance, we want to compare how the latents and raw curves perform on different size data sets. We take our initial dataset, and split it into 80% training data and 20% test data. From there, we subset the training data to create smaller training datasets to see how the models perform with less training data. For each type of input, we train a model to predict either the specific value (regression) or category (classification). We then compare the accuracy of the models on a test dataset withheld from training.

### Simulated consortia data generation and preprocessing for testing ML-based forecasting

150 consortia were simulated using the chaotic consortia model [30] and the bounded generalized Lotka-Volterra model [33] [Supp Fig 9A]. The consortia were generated with a total number of members, and from that a subset were selected as focal members [Supp Fig 9B]. Each consortia had 10,000 simulations. In all of the simulations the focal community initial cell densities were randomized. In some of the simulations the background strain starting cell densities were randomized, but in the remaining the starting cell densities were fixed. Using the maximum likelihood estimator [7], we estimated the intrinsic dimensions for each of the consortia and selected 2 for our analysis: a simple consortium with 3 estimated intrinsic dimensions and a complex consortium with 6 estimated intrinsic dimensions [Supp Fig 9C]. The simple community had 5 focal members with 15 background strains, and the data were generated using the bounded generalized Lotka-Volterra model. The complex community had 8 focal members with 92 background strains, and the data were generated using the chaotic consortia model.

### Experimental consortia data preprocessing for testing ML-based forecasting

We used a dataset generated by Fujita *et al.* to predict consortia behavior over time [4]. The dataset had 3 different media conditions and 2 different source microbial consortia, resulting in 6 different experiment conditions. For each of the 6 experiment conditions, they ran 8 replicates. They took daily readings for 110 days, and each of the experiment conditions had anywhere from 28 to 144 strains. However, not all the strains were present in each of the replicates, so for each experiment condition we selected a focal community consisting of the strains that were present across all 8 replicates. This resulted in focal communities ranging from 2 to 18 members [Supp]. For each of the 6 experiment conditions, we performed eightfold cross-validation to train our models. With each fold, one of the replicates was used as a test dataset and the training was completed on the other 7 replicates. Due to the fact this data is daily readings, we interpolate the data. We linearly interpolated the 110 time points to 1,024 time points, resulting in approximately 13.75 days of data per 128 time points.

### Consortia prediction and forecasting using ML

The pipeline for consortia prediction and forecasting was the same for both the simulations and the experimental data. We generated datasets by taking 128 time point windows from the data and predicting the next 128 time points. We predicted all focal members of the consortia simultaneously, so the input and output vectors were of shape *M* number of members x 128 time points. For each member, the 128 time point window is encoded into its latent representation, and we include the maximum value from the window. This results in input and output vectors of shape *M* x 9.

We trained two separate regression models: one using the raw curves and one using the latent vectors. For the raw curves, we flattened the input and output to change them from shape *M* x 128 to 128*M* x 1. For the latent vectors, we flattened the input and output to change them from shape *M* x 9 to 9*M* x 1. We then used these flattened vectors to train the regression models.

To forecast into the future, we used the trained regression model. We begin with a seed of the first 128 time points from a simulation. Then, we iteratively predicted the next 128-time-point chunk and slid the window of the input. This allows us to forecast indefinitely into the future from an initial seed.

Each consortia was individually trained on its own regression models. To characterize the forecast accuracy of the model we consider the root mean square error (RMSE) over time. RMSE is used for the forecasting task, because its units match the underlying consortia. It is considered over time, because this allows us to analyze for how long the predictions are accurate.

### Consortia final abundance classification using ML

For the experimental consortia, we wanted to predict whether the final abundance for each of the members would be above or below 0.1. To do this, we took the first 128 time points, which equated to approximately 14 days, and trained a classifier to predict whether each member of the consortium would be above or below 0.1 in the final time point, 96 days later. Similar to the forecasting, we predicted all members of the consortia simultaneously, so the input vectors were of shape *M* number of members x 128 time points and the binary output vectors were of shape *M* x 1. For each member, the 128 time point window is encoded into its latent representation, and we include the maximum value from the window. This results in input and output vectors of shape *M* x 9.

We trained two separate classification models: one using the raw curves and one using the latent vectors. For the raw curves, we flattened the input to change them from shape *M* x 128 to 128*M* x 1. For the latent vectors, we flattened the input to change them from shape *M* x 9 to 9*M* x 1. We then used these flattened vectors to train the classification models.

Each consortia was individually trained on its own classification models. We used classification accuracy to report the results, which reports the fraction of the samples that were correctly classified.

## Acknowledgements

This work was supported by National Institute of Health (L.Y.: R01AI125604, R01GM098642, R01EB031869), DARPA (L.Y. HR0011-23-2-0008), and the National Science Foundation (L. Y.: Cooperative Agreement No. EEC-2133504, and a graduate fellowship to Z. A. H.).

## Author contributions

Z.A.H. and L.Y. conceived the research and designed the research framework. Z.A.H. and I.S. performed all computation, model simulation, neural network training, and machine learning analysis with input from A.L.H., X.W., P.C., and, L.Y.. Z.A.H., A.L.H., K.E.D., H.R.M., D.L., R.M., K.Kim, E.Ş., G.S.H., H.S., C.A.V., J.L., Y.H., A.R.S., Z.Y., S.L., D.M.S., H.D., F.W., S.W., and E.C. generated experimental data. M.L., A.J.L., and L.D. provided experimental data. Z.Z. generated numerical simulations for microbial consortia tasks. K.Kholina developed the transformer-based model with input from Z.A.H. and P.C.. Z.A.H., I.S., and L.Y. wrote the manuscript with input from H.R.M., Z.Z., D.L., R.M., E.Ş., J.L., Y.B., and S.W..

## Data availability

All data are available in the main text, supplemental, in our GitHub repository (https://github.com/youlab/foundation), or in our Hugging Face pages for models (https://huggingface.co/you-lab/models) and datasets (https://huggingface.co/you-lab/datasets). Instructions for use of the Hugging Face collection are included in the GitHub README files.

## Code availability

All code for the project, including data collection, model training, and figure generation, is available in our GitHub repository (https://github.com/youlab/foundation).

## References

[1] C. Zhang et al., “Temporal encoding of bacterial identity and traits in growth dynamics,” Proc. Natl. Acad. Sci., vol. 117, no. 33, pp. 20202–20210, Aug. 2020, doi: 10.1073/pnas.2008807117.

[2] Y. Ram et al., “Predicting microbial growth in a mixed culture from growth curve data,” Proc. Natl. Acad. Sci., vol. 116, no. 29, pp. 14698–14707, July 2019, doi: 10.1073/pnas.1902217116.

[3] M. Baranwal, R. L. Clark, J. Thompson, Z. Sun, A. O. Hero, and O. S. Venturelli, “Recurrent neural networks enable design of multifunctional synthetic human gut microbiome dynamics,” eLife, vol. 11, p. e73870, June 2022, doi: 10.7554/eLife.73870.

[4] H. Fujita et al., “Alternative stable states, nonlinear behavior, and predictability of microbiome dynamics,” Microbiome, vol. 11, no. 1, Art. no. 1, Dec. 2023, doi: 10.1186/s40168-023-01474-5.

[5] G. Armstrong et al., “Applications and Comparison of Dimensionality Reduction Methods for Microbiome Data,” Front. Bioinforma., vol. 2, Feb. 2022, doi: 10.3389/fbinf.2022.821861.

[6] M. A. Carreira-Perpinan, “A Review of Dimension Reduction Techniques”.

[7] E. Levina and P. J. Bickel, “Maximum Likelihood Estimation of Intrinsic Dimension,” in Advances in Neural Information Processing Systems, MIT Press, 2004. [Online]. Available: https://papers.nips.cc/paper_files/paper/2004/hash/74934548253bcab8490ebd74afed7031-Abstract.html

[8] Y. Baig, H. R. Ma, H. Xu, and L. You, “Autoencoder neural networks enable low dimensional structure analyses of microbial growth dynamics,” Nat. Commun., vol. 14, no. 1, p. 7937, Dec. 2023, doi: 10.1038/s41467-023-43455-0.

[9] Z. Zhou, et al., “Linear scaling reveals low-dimensional structure in observable microbial dynamics,” June 19, 2025, bioRxiv. doi: 10.1101/2025.06.13.659614.

[10] “Simulating 500 million years of evolution with a language model | Science.” Accessed: Feb. 27, 2025. [Online]. Available: https://www-science-org.proxy.lib.duke.edu/doi/10.1126/science.ads0018

[11] E. Nguyen et al., “Sequence modeling and design from molecular to genome scale with Evo,” Science, vol. 386, no. 6723, p. eado9336, Nov. 2024, doi: 10.1126/science.ado9336.

[12] “scBERT as a large-scale pretrained deep language model for cell type annotation of single-cell RNA-seq data | Nature Machine Intelligence.” Accessed: Jan. 21, 2025. [Online]. Available: https://www.nature.com/articles/s42256-022-00534-z

[13] A. B. George and K. S. Korolev, “Ecological landscapes guide the assembly of optimal microbial communities,” PLOS Comput. Biol., vol. 19, no. 1, p. e1010570, Jan. 2023, doi: 10.1371/journal.pcbi.1010570.

[14] S. Arya, A. B. George, and J. P. O’Dwyer, “Sparsity of higher-order landscape interactions enables learning and prediction for microbiomes,” Proc. Natl. Acad. Sci., vol. 120, no. 48, p. e2307313120, Nov. 2023, doi: 10.1073/pnas.2307313120.

[15] “Enabling high-throughput biology with flexible open-source automation | Molecular Systems Biology.” Accessed: Jan. 22, 2025. [Online]. Available: https://www.embopress.org/doi/full/10.15252/msb.20209942

[16] H. R. Ma, H. Z. Xu, K. Kim, D. J. Anderson, and L. You, “Private benefit of β-lactamase dictates selection dynamics of combination antibiotic treatment,” Nat. Commun., vol. 15, no. 1, p. 8337, Sept. 2024, doi: 10.1038/s41467-024-52711-w.

[17] J. N. Hennigan, R. Menacho-Melgar, P. Sarkar, M. Golovsky, and M. D. Lynch, “Scalable, robust, high-throughput expression & purification of nanobodies enabled by 2-stage dynamic control,” Metab. Eng., vol. 85, pp. 116–130, Sept. 2024, doi: 10.1016/j.ymben.2024.07.012.

[18] S. Li, Z. Ye, E. A. Moreb, R. Menacho-Melgar, M. Golovsky, and M. D. Lynch, “2-Stage microfermentations,” Metab. Eng. Commun., vol. 18, p. e00233, June 2024, doi: 10.1016/j.mec.2024.e00233.

[19] S. Li et al., “Dynamic control over feedback regulatory mechanisms improves NADPH flux and xylitol biosynthesis in engineered *E. coli*,” Metab. Eng., vol. 64, pp. 26–40, Mar. 2021, doi: 10.1016/j.ymben.2021.01.005.

[20] R. Menacho-Melgar et al., “Scalable, two-stage, autoinduction of recombinant protein expression in E. coli utilizing phosphate depletion,” Biotechnol. Bioeng., vol. 117, no. 9, pp. 2715–2727, 2020, doi: 10.1002/bit.27440.

[21] M. Ahmad et al., “Tradeoff between lag time and growth rate drives the plasmid acquisition cost,” Nat. Commun., vol. 14, no. 1, p. 2343, Apr. 2023, doi: 10.1038/s41467-023-38022-6.

[22] S. V. Aduru et al., “Sub-inhibitory antibiotic treatment selects for enhanced metabolic efficiency,” Microbiol. Spectr., vol. 12, no. 2, pp. e03241–23, Jan. 2024, doi: 10.1128/spectrum.03241-23.

[23] A. Palomino et al., “Metabolic genes on conjugative plasmids are highly prevalent in Escherichia coli and can protect against antibiotic treatment,” ISME J., vol. 17, no. 1, pp. 151–162, Jan. 2023, doi: 10.1038/s41396-022-01329-1.

[24] H. Prensky, A. Gomez-Simmonds, A. Uhlemann, and A. J. Lopatkin, “Conjugation dynamics depend on both the plasmid acquisition cost and the fitness cost,” Mol. Syst. Biol., vol. 17, no. 3, p. e9913, Mar. 2021, doi: 10.15252/msb.20209913.

[25] R. Maddamsetti et al., “Duplicated antibiotic resistance genes reveal ongoing selection and horizontal gene transfer in bacteria,” Nat. Commun., vol. 15, no. 1, p. 1449, Feb. 2024, doi: 10.1038/s41467-024-45638-9.

[26] Z. Ye et al., “Two-stage dynamic deregulation of metabolism improves process robustness & scalability in engineered *E. coli*.,” Metab. Eng., vol. 68, pp. 106–118, Nov. 2021, doi: 10.1016/j.ymben.2021.09.009.

[27] L. A. David et al., “Host lifestyle affects human microbiota on daily timescales,” Genome Biol., vol. 15, no. 7, p. R89, July 2014, doi: 10.1186/gb-2014-15-7-r89.

[28] J. Schluter et al., “The gut microbiota is associated with immune cell dynamics in humans,” Nature, vol. 588, no. 7837, pp. 303–307, Dec. 2020, doi: 10.1038/s41586-020-2971-8.

[29] C. V. Theodoris et al., “Transfer learning enables predictions in network biology,” Nature, vol. 618, no. 7965, pp. 616–624, June 2023, doi: 10.1038/s41586-023-06139-9.

[30] J. Hu, D. R. Amor, M. Barbier, G. Bunin, and J. Gore, “Emergent phases of ecological diversity and dynamics mapped in microcosms,” Science, vol. 378, no. 6615, pp. 85–89, Oct. 2022, doi: 10.1126/science.abm7841.

[31] F. Yang, F. Wang, L. Huang, L. Liu, J. Huang, and J. Yao, “Reply to: Deeper evaluation of a single-cell foundation model,” *Nat*. Mach. Intell., vol. 6, no. 12, pp. 1447–1450, Dec. 2024, doi: 10.1038/s42256-024-00948-x.

[32] J. E. Sulaiman et al., “Elucidating human gut microbiota interactions that robustly inhibit diverse Clostridioides difficile strains across different nutrient landscapes,” Nat. Commun., vol. 15, no. 1, p. 7416, Aug. 2024, doi: 10.1038/s41467-024-51062-w.

[33] F. Wu et al., “Modulation of microbial community dynamics by spatial partitioning,” Nat. Chem. Biol., vol. 18, no. 4, pp. 394–402, Apr. 2022, doi: 10.1038/s41589-021-00961-w.

[34] R. Lam et al., “Learning skillful medium-range global weather forecasting,” Science, vol. 382, no. 6677, pp. 1416–1421, Dec. 2023, doi: 10.1126/science.adi2336.

[35] “Complementing 16S rRNA Gene Amplicon Sequencing with Total Bacterial Load To Infer Absolute Species Concentrations in the Vaginal Microbiome | mSystems.” Accessed: Oct. 21, 2025. [Online]. Available: https://journals.asm.org/doi/10.1128/msystems.00777-19

[36] G. B. Gloor, J. M. Macklaim, V. Pawlowsky-Glahn, and J. J. Egozcue, “Microbiome Datasets Are Compositional: And This Is Not Optional,” Front. Microbiol., vol. 8, Nov. 2017, doi: 10.3389/fmicb.2017.02224.

[37] J. T. Barlow, S. R. Bogatyrev, and R. F. Ismagilov, “A quantitative sequencing framework for absolute abundance measurements of mucosal and lumenal microbial communities,” Nat. Commun., vol. 11, no. 1, p. 2590, May 2020, doi: 10.1038/s41467-020-16224-6.

[38] “Metagenomic estimation of absolute bacterial biomass in the mammalian gut through host-derived read normalization | mSystems.” Accessed: Oct. 21, 2025. [Online]. Available: https://journals.asm.org/doi/10.1128/msystems.00984-25

[39] “Enabling high-throughput biology with flexible open-source automation | Molecular Systems Biology.” Accessed: Jan. 09, 2025. [Online]. Available: https://www.embopress.org/doi/full/10.15252/msb.20209942

[40] T. Baba et al., “Construction of Escherichia coli K-12 in-frame, single-gene knockout mutants: the Keio collection,” Mol. Syst. Biol., vol. 2, no. 1, p. 2006.0008, Feb. 2006, doi: 10.1038/msb4100050.

[41] T. Akiba, S. Sano, T. Yanase, T. Ohta, and M. Koyama, “Optuna: A Next-generation Hyperparameter Optimization Framework,” in Proceedings of the 25th ACM SIGKDD International Conference on Knowledge Discovery & Data Mining, in KDD ’19. New York, NY, USA: Association for Computing Machinery, July 2019, pp. 2623–2631. doi: 10.1145/3292500.3330701.

[42] F. Pedregosa et al., “Scikit-learn: Machine Learning in Python,” J. Mach. Learn. Res., vol. 12, no. 85, pp. 2825–2830, 2011.

