## Supplemental Information for "A foundation model for microbial growth dynamics"

Current affiliations

✳ Department of Molecular, Cellular, and Developmental Biology, Yale University, New Haven, CT, USA

† Department of Biochemistry and Microbiology, Rutgers University, New Brunswick, NJ, USA

¶ Department of Physics, University of Florida, Gainesville, FL, USA

‡ Department of Systems, Synthetic, and Physical Biology, Rice University, Houston, TX, USA

✢ Department of Bioengineering, Stanford University, Stanford, CA, USA

§ The Ensynble, Inc., San Francisco, CA, USA

‖ Bay Area Institute of Computation, Altos Labs, Redwood City, CA, USA.

### Supplemental

#### 1. Model descriptions

All of our models are designed to take 128 time point vectors as inputs and encode the input to a smaller dimension, ranging from 2 to 24. The models were trained using a loss function of mean squared error, with the additional Kullbeck-Leibler divergence loss for the variational autoencoders. The training was run for a given number of epochs, but the training was stopped early when the test loss stopped improving. Given the multitude of possible parameters in terms of number of layers, learning rates, channels, etc., we first ran optimizations to pick the best values for each model type. These optimizations were completed either by manually testing different configurations or using Optuna to optimize the test loss of the model [1]. The following descriptions of the models are within the previously described framework.

**Autoencoder7X (A7X)** is a custom variational autoencoder with convolution and linear layers for processing microbial growth curve data implemented in PyTorch [2]. The encoder consists of 7 layers. The first 6 are convolution layers, which use a combination of kernel sizes, strides, dilations, and padding to either keep the vector flat or reduce the length by 2 while altering the number of channels. The final layer of the encoder is the multi-layer perceptron (MLP) layer. The MLP layer takes the output of the last convolution layer and reduces it to the latent dimension specified in the initialization of the model. During training, there are two MLP layers: mu and logvar. The mu is the mean value for the vector, and the logvar applies random noise to the mean such that each input corresponds to a continuous latent range, and not a single value. During evaluation, only the mu layer is used. The decoder consists of 8 layers. The first layer is a linear layer that expands the vector, which is then input into 7 consecutive transposed convolution layers. Similar to the encoder, the transposed convolution layers either keep the vector flat or expand by 2 while adjusting the channels. With the exception of the layer preceding the latent vector and the reconstructed output, each layer is followed by the rectified linear unit activation function. This activation function sets all values below 0 to 0.

**Multi-layer perceptron Network Model (MNM)** is a custom variational autoencoder consisting solely of linear layers implemented in PyTorch [2]. The encoder is a multi-layer perceptron (MLP) with a given number of layers and hidden dimensions. The output of the MLP is then fed into the mu and logvar layers, as described in Autoencoder7X. The decoder then takes the latent vector into another MLP, and the output is the reconstructed growth curve. With the exception of the layer preceding the latent vector and the reconstructed output, each layer is followed by the rectified linear unit activation function. This activation function sets all values below 0 to 0.

**Variational autoencoder Bottleneck (VB)** is a custom variational autoencoder consisting solely of convolution layers implemented in PyTorch [2]. Unlike the previous two models which use linear layers to specify the latent size, this model uses the kernel size and stride of the convolution layers to encode and then decode the latent vector. The encoder consists of 7 layers, including a final layer with mu and logvar layers which are convolution layers. The decoder consists of 7 transposed convolution layers. With the exception of the layer preceding the latent vector and the reconstructed output, each layer is followed by the rectified linear unit activation function. This activation function sets all values below 0 to 0.

**MicroBERT Curve Reducer (MCR)** is a custom BERT-based model designed for processing microbial growth curve data implemented in PyTorch [2]. The processing pipeline begins with input masking, where 15% of the vector values are randomly masked (replaced with 0). The masked input is then projected through a linear layer from its original 1x128 dimension to a smaller hidden dimension. This compressed representation passes through transformer encoder layers with attention heads. The final MLM (Masked Language Model) head, implemented as a linear layer, converts the data back to the original dimension. During training, the mean squared error loss is computed only between the predicted and actual values at masked positions. The model is trained with the learning rate of 1E-2, 1E-3, or 1E-4. The training is run for a maximum of 10,000 epochs, but the training is stopped once the test loss does not improve for 20 consecutive epochs.

**PCA Reducer (PR)** is a standard implementation of principal component analysis (PCA) using the scikit-learn library [3]. PCA is a common dimension reduction technique used broadly in many different areas [4], [5]. We use the IncrementalPCA implementation due to the size of our dataset. The class has built-in methods to transform input data to the defined latent vector size, and to perform inverse transformation of the latent vectors, which we use for the reconstruction task.

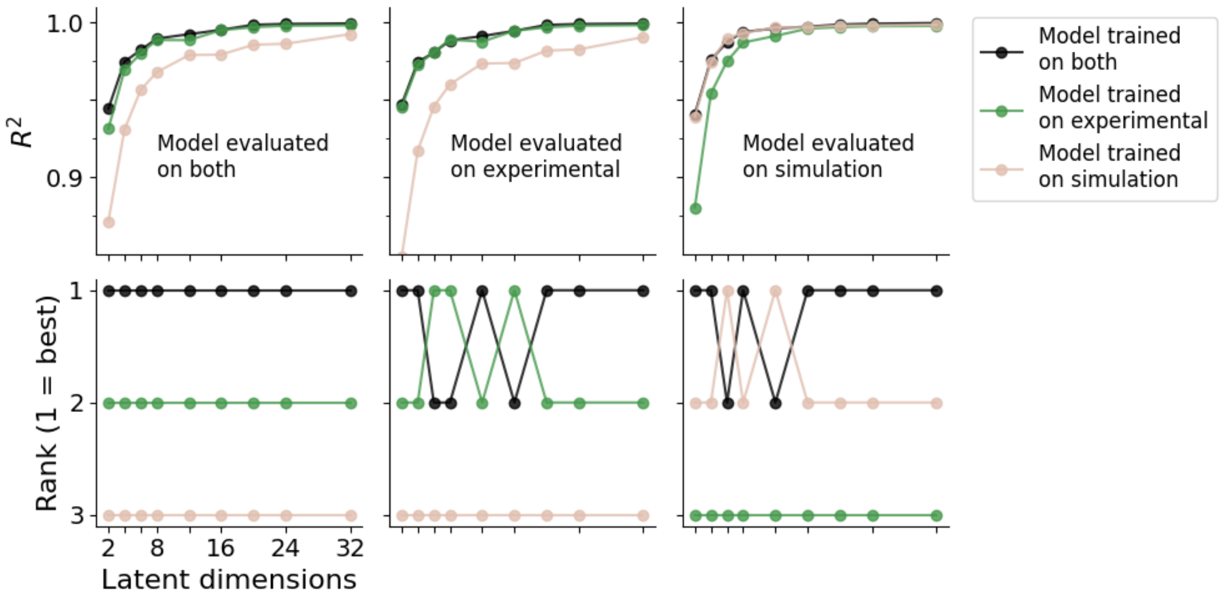


**Supplemental Figure S1. At most latent dimensions, the reconstruction accuracy is higher on the experimental dataset when the foundation model is trained on both experimental and simulation data**

We trained 27 different versions of our custom variational autoencoder, Autoencoder7X (A7X). The 27 models were a combination of 3 training dataset types and 9 latent dimension sizes. The 3 training datasets were only experimental data, only simulation data, and both experimental and simulation data. The latent sizes were 2, 4, 6, 8, 12, 16, 20, 24, and 32. The models trained on both datasets outperformed the other two models at all latent sizes. The models trained on both datasets outperformed the models trained on experimental data when evaluated on only the experimental data at 6 of the 9 latent dimension sizes. The models trained on both datasets outperformed the models trained on simulation data when evaluated on only the simulation data at 7 of the 9 latent dimension sizes.

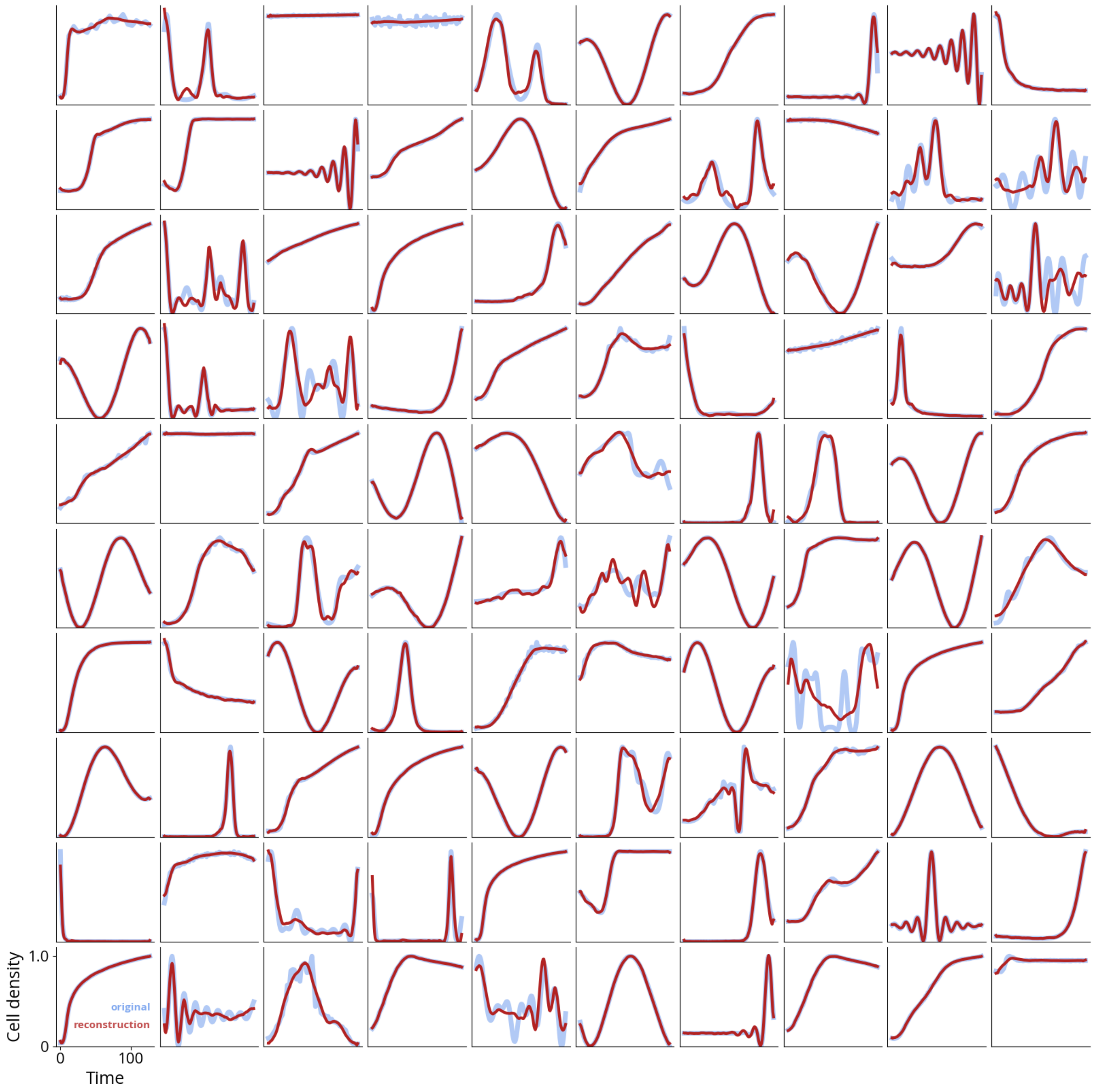


**Supplemental Figure S2. Model reconstructions: 100 randomly selected samples from the test dataset for the model training.**

Using our test dataset from model training, we randomly selected 100 growth curves to reconstruct using the Autoencoder7X model with 8 latent dimensions. The model is able to accurately reconstruct simple curves, and still provides meaningful reconstruction on more complex curves.

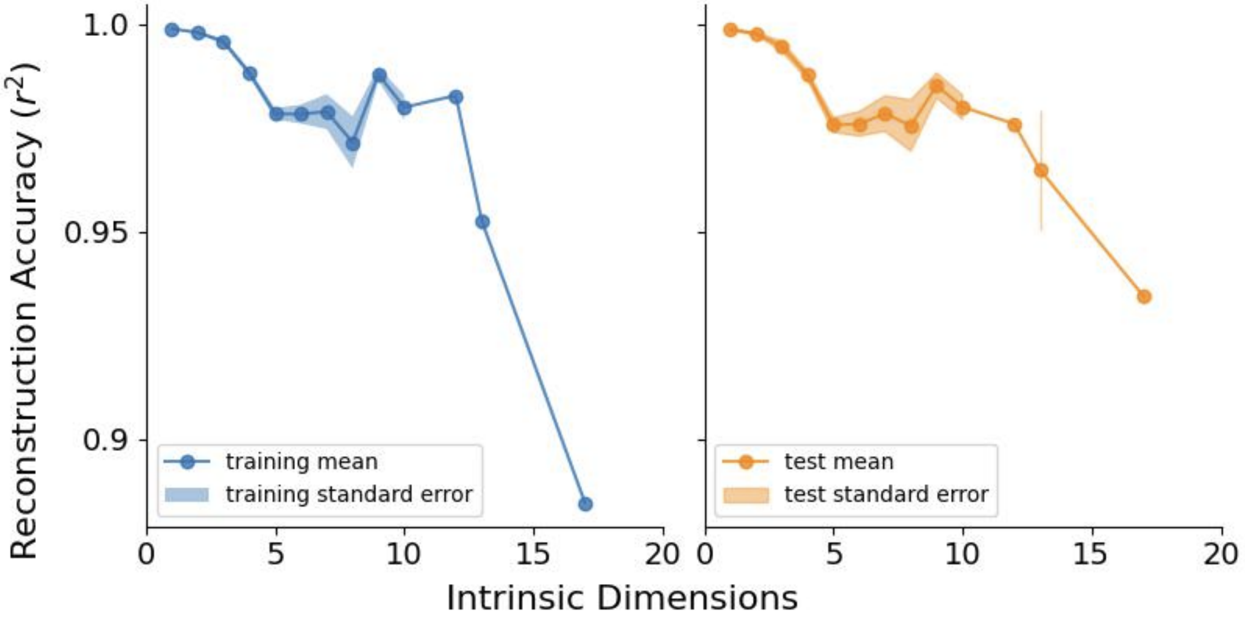


**Supplemental Figure S3. Reconstruction accuracy is higher on datasets with lower estimated intrinsic dimensions**

There were 1,225 unique data files used for this study. For each of the data files, we estimated the intrinsic dimensions of the dataset. Then, we reconstructed the data using the Autoencoder7X model with 8 latent dimensions. We see that the reconstruction accuracy is better on datasets with lower intrinsic dimensions, and worse on datasets with higher intrinsic dimensions.

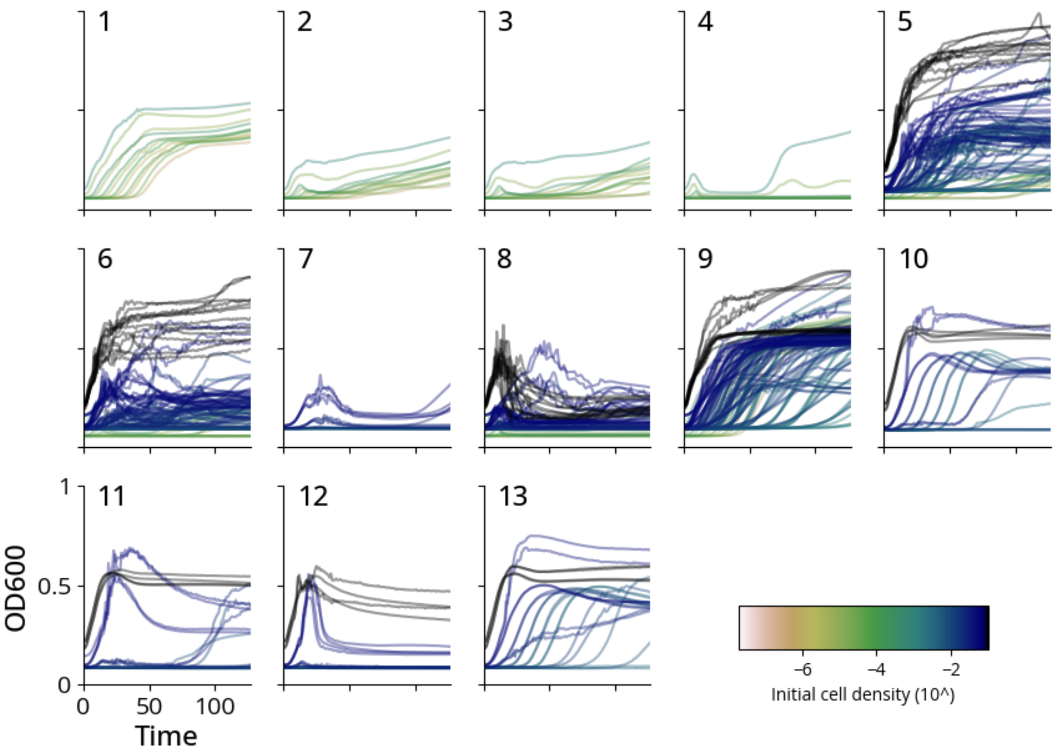


**Supplemental Figure S4. Antibiotic dataset for treatment and concentration featuring TOP10F at different initial cell densities**

672 growth curves from experiments completed using TOP10F. Initial cell densities were varied, as indicated by the color scale. Experimental conditions used (media, antibiotics, concentration (μg/mL)):

1. LB, amoxicillin 1.0
2. LB amoxicillin 2.0
3. LB amoxicillin 2.2
4. LB amoxicillin 4.0
5. LB carbenicillin 5.0
6. LB carbenicillin 10.0
7. LB carbenicillin 15.0

8. LB carbenicillin 30.0
9. LB none 0.0
10. M9CA carbenicillin 5.0
11. M9CA carbenicillin 10.0
12. M9CA carbenicillin 30.0
13. M9CA none 0.0

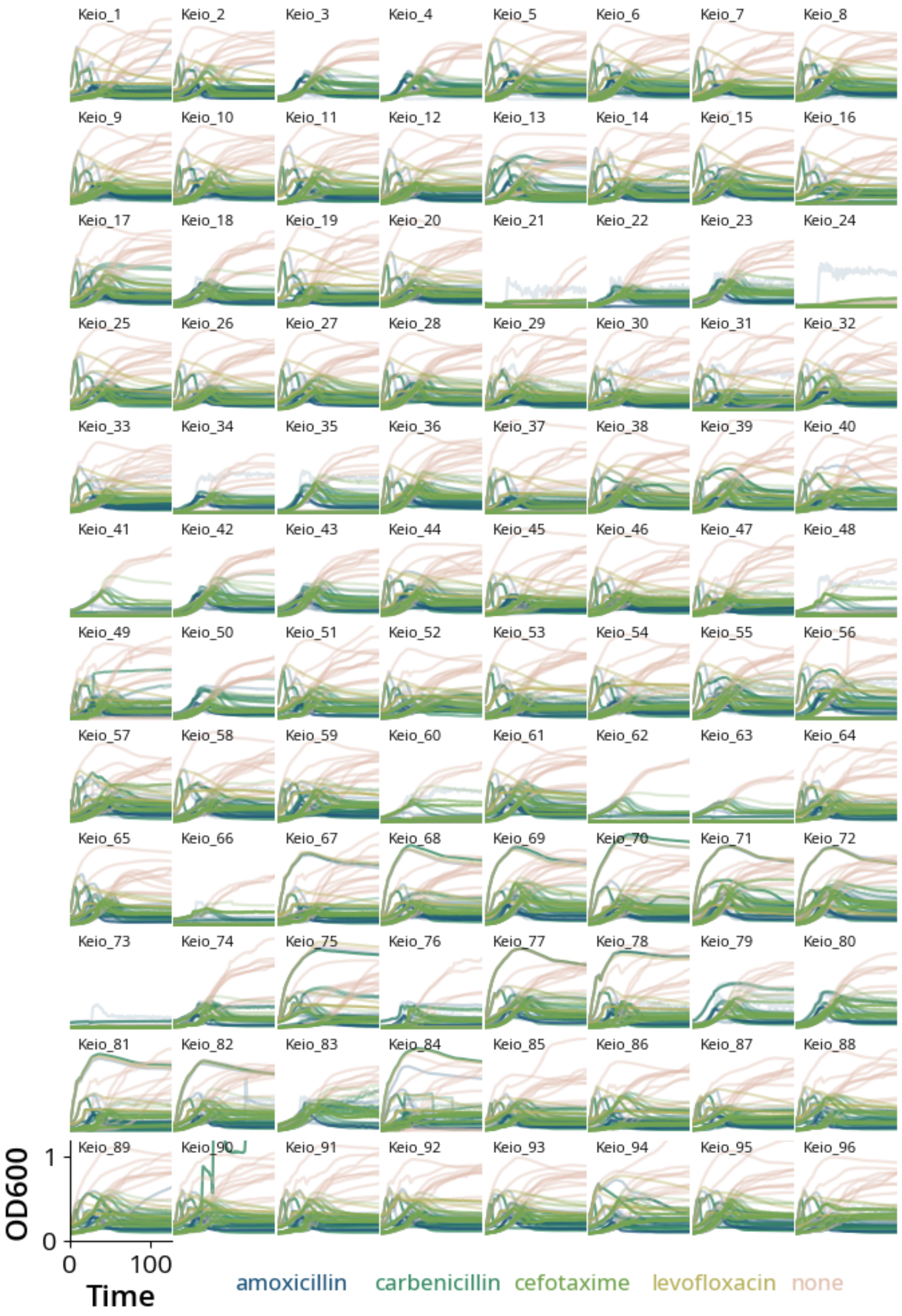
Keio_1
Keio_2
Keio_3
Keio_4
Keio_5
Keio_6
Keio_7
Keio_8
Keio_9
Keio_10
Keio_11
Keio_12
Keio_13
Keio_14
Keio_15
Keio_16
Keio_17
Keio_18
Keio_19
Keio_20
Keio_21
Keio_22
Keio_23
Keio_24
Keio_25
Keio_26
Keio_27
Keio_28
Keio_29
Keio_30
Keio_31
Keio_32
Keio_33
Keio_34
Keio_35
Keio_36
Keio_37
Keio_38
Keio_39
Keio_40
Keio_41
Keio_42
Keio_43
Keio_44
Keio_45
Keio_46
Keio_47
Keio_48
Keio_49
Keio_50
Keio_51
Keio_52
Keio_53
Keio_54
Keio_55
Keio_56
Keio_57
Keio_58
Keio_59
Keio_60
Keio_61
Keio_62
Keio_63
Keio_64
Keio_65
Keio_66
Keio_67
Keio_68
Keio_69
Keio_70
Keio_71
Keio_72
Keio_73
Keio_74
Keio_75
Keio_76
Keio_77
Keio_78
Keio_79
Keio_80
Keio_81
Keio_82
Keio_83
Keio_84
Keio_85
Keio_86
Keio_87
Keio_88
Keio_89
Keio_90
Keio_91
Keio_92
Keio_93
Keio_94
Keio_95
Keio_96
OD600
1
0
0
100
Time
amoxicillin
carbenicillin
cefotaxime
levofloxacin
none

**Supplemental Figure S5. Antibiotic dataset for treatment and concentration featuring Keio strains**

5,100 growth curves from experiments completed using the Keio strains. There were 4 different antibiotics used, as indicated by colors. The antibiotics were applied at different concentrations, with darker lines indicating higher antibiotic concentration. For specific experimental parameters, reference data files.

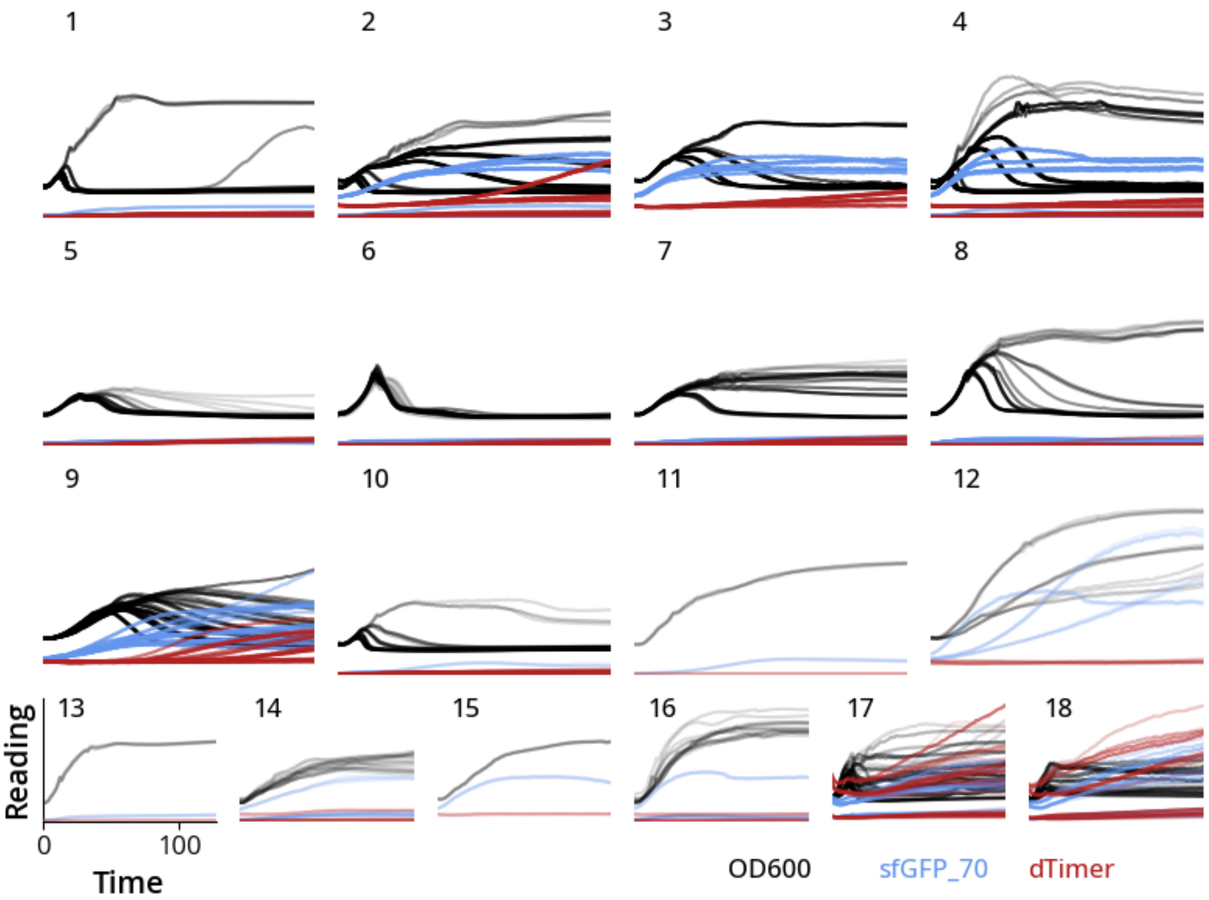


**Supplemental Figure S6. Antibiotic dataset for treatment and concentration featuring MG1655 with fluorescent plasmid**

1,152 growth curves from experiments completed using *E. coli* MG1655 with p15A-pTet-sfGFP-linker-Tdimer-kanR plasmid. The antibiotics were applied at different concentrations, with darker lines indicating higher antibiotic concentration. For specific experimental parameters, reference data files. There were 18 different experimental conditions (media, antibiotic, casamino acid %):

1. M9CA+0.4%glucose, carbenicillin, 0.0
2. M9CA+0.4%glucose, carbenicillin, 0.02
3. M9CA+0.4%glucose, carbenicillin, 0.1
4. M9CA+0.4%glucose, carbenicillin, 0.2
5. M9CA+0.4%glucose, cefotaxime, 0.02
6. M9CA+0.4%glucose, cefotaxime, 0.2
7. M9CA+0.4%glucose, amoxicillin, 0.02
8. M9CA+0.4%glucose, amoxicillin, 0.2,

9. M9CA+0.4%glucose, carbenicillin/chloramphenicol, 0.2
10. M9CA+0.4%glucose, carbenicillin/chloramphenicol, 0.0,
11. M9CA+0.4%glucose, chloramphenicol, 0.0
12. M9CA+0.4%glucose, chloramphenicol, 0.2
13. M9CA+0.4%glucose, none, 0.0
14. M9CA+0.4%glucose, none, 0.02
15. M9CA+0.4%glucose, none, 0.1
16. M9CA+0.4%glucose, none, 0.2
17. LB, none, 0.0
18. tbroth, none, 0.0

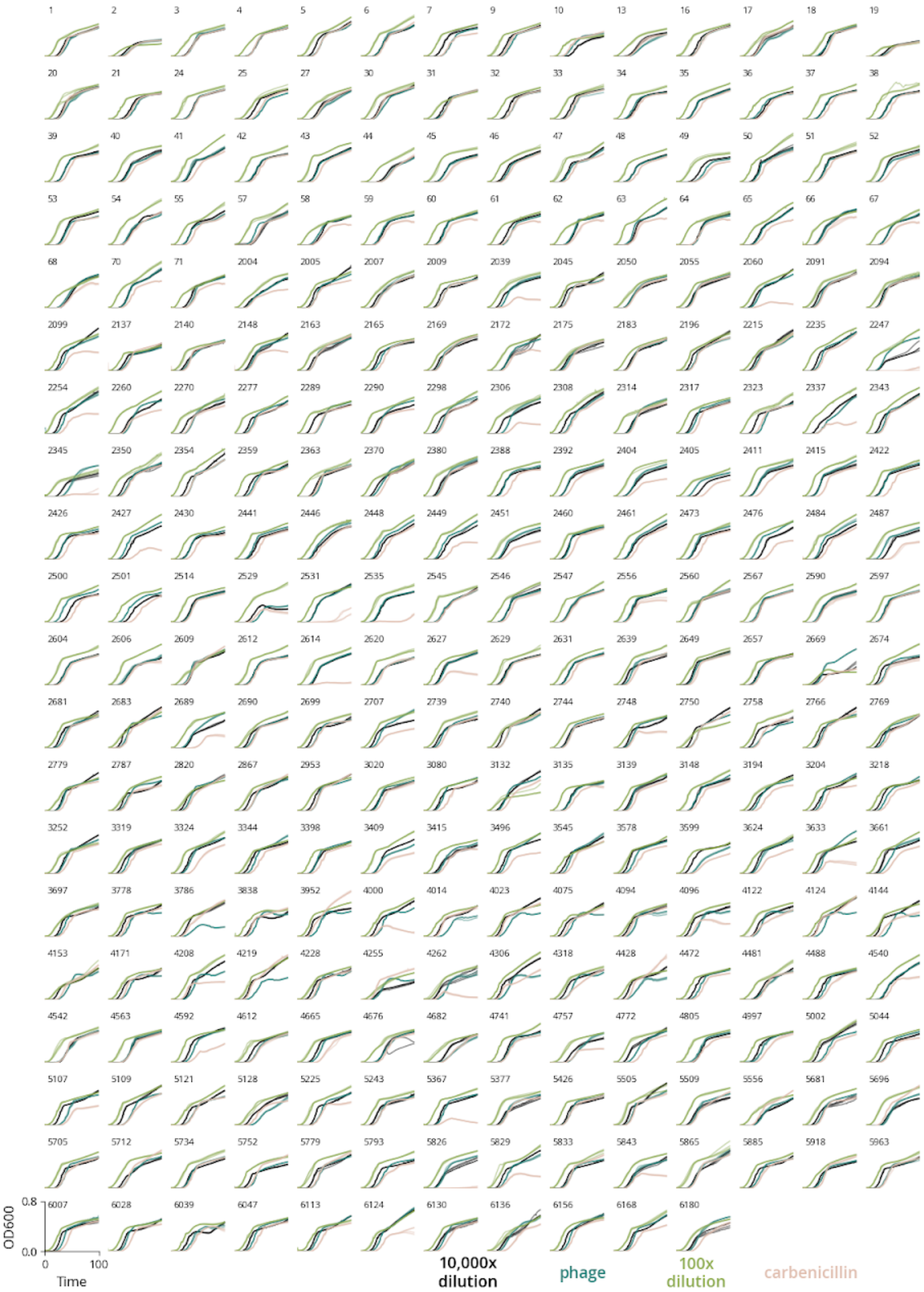

1 2 3 4 5 6 7 9 10 13 16 17 18 19
20 21 24 25 27 30 31 32 33 34 35 36 37 38
39 40 41 42 43 44 45 46 47 48 49 50 51 52
53 54 55 57 58 59 60 61 62 63 64 65 66 67
68 70 71 2004 2005 2007 2009 2039 2045 2050 2055 2060 2091 2094
2099 2137 2140 2148 2163 2165 2169 2172 2175 2183 2196 2215 2235 2247
2254 2260 2270 2277 2289 2290 2298 2306 2308 2314 2317 2323 2337 2343
2345 2350 2354 2359 2363 2370 2380 2388 2392 2404 2405 2411 2415 2422
2426 2427 2430 2441 2446 2448 2449 2451 2460 2461 2473 2476 2484 2487
2500 2501 2514 2529 2531 2535 2545 2546 2547 2556 2560 2567 2590 2597
2604 2606 2609 2612 2614 2620 2627 2629 2631 2639 2649 2657 2669 2674
2681 2683 2689 2690 2699 2707 2739 2740 2744 2748 2750 2758 2766 2769
2779 2787 2820 2867 2953 3020 3080 3132 3135 3139 3148 3194 3204 3218
3252 3319 3324 3344 3398 3409 3415 3496 3545 3578 3599 3624 3633 3661
3697 3778 3786 3838 3952 4000 4014 4023 4075 4094 4096 4122 4124 4144
4153 4171 4208 4219 4228 4255 4262 4306 4318 4428 4472 4481 4488 4540
4542 4563 4592 4612 4665 4676 4682 4741 4757 4772 4805 4997 5002 5044
5107 5109 5121 5128 5225 5243 5367 5377 5426 5505 5509 5556 5681 5696
5705 5712 5734 5752 5779 5793 5826 5829 5833 5843 5865 5885 5918 5963
6007 6028 6039 6047 6113 6124 6130 6136 6156 6168 6180
0.8
0.0
OD600
0
100
Time
10,000x dilution
phage
100x dilution
carbenicillin

**Supplemental Figure S7. Antibiotic dataset for resistance**

4,432 growth curves from the experiment using 277 different clinical isolates. These isolates were collected from Duke Hospital, and they are of the genuses *Klebsiella, Citrobacter,* and *Escherichia*. Of the 277 strains, 42 are resistant to none of the antibiotics, 47 are resistant to 1, 114 are resistant to 2, 45 are resistant to 3, and 29 are resistant to all 4. Each panel is the 16 growth curves from each isolate. The strains were grown in 4 different media conditions, and each condition had 4 replicates. These 4 different conditions are as follows: 10,000x dilution from overnight sample, addition of λ phage, 100x dilution from overnight sample, 5 μg/mL carbenicillin. For specifics on the experimental protocol, reference methods from Zhang *et al.* [6].

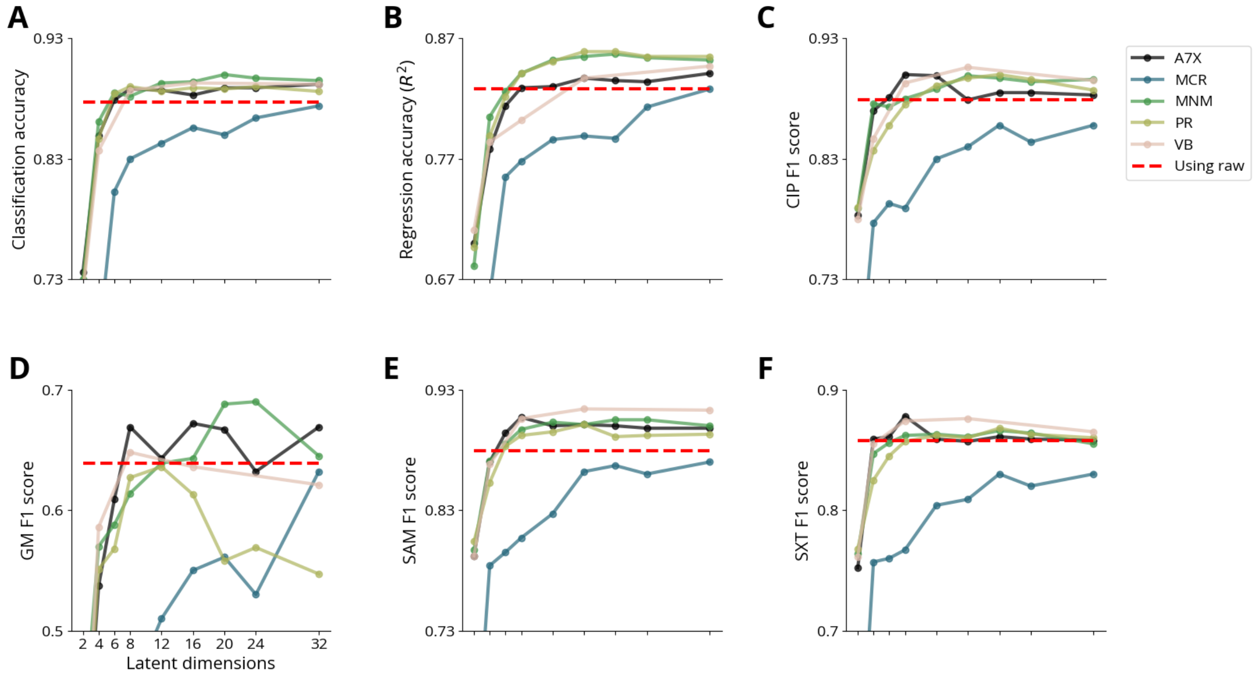


**Supplemental Figure S8. Summary of the performance of the different models on the antibiotic applications**

For all of the antibiotic tasks from the main text, we wanted to compare the results of our different models. For each task, we took growth curves as inputs, and converted the growth curves to latent vectors. We then used these latent vectors as the input into a regression or classification model.

A) Classification accuracy of antibiotic type, reported as samples correctly classified. This is the task presented in Fig 3A.

B) Prediction accuracy of antibiotic concentration, reported as regression accuracy ($R^2$). This is the task presented in Fig 3C.

C) Classification accuracy of CIP resistance using each model, reported as F1 score. This is the task presented in Fig 3E.

D) Classification accuracy of GM resistance using each model, reported as F1 score. This is the task presented in Fig 3E.

E) Classification accuracy of SAM resistance using each model, reported as F1 score. This is the task presented in Fig 3E.

F) Classification accuracy of SXT resistance using each model, reported as F1 score. This is the task presented in Fig 3E.

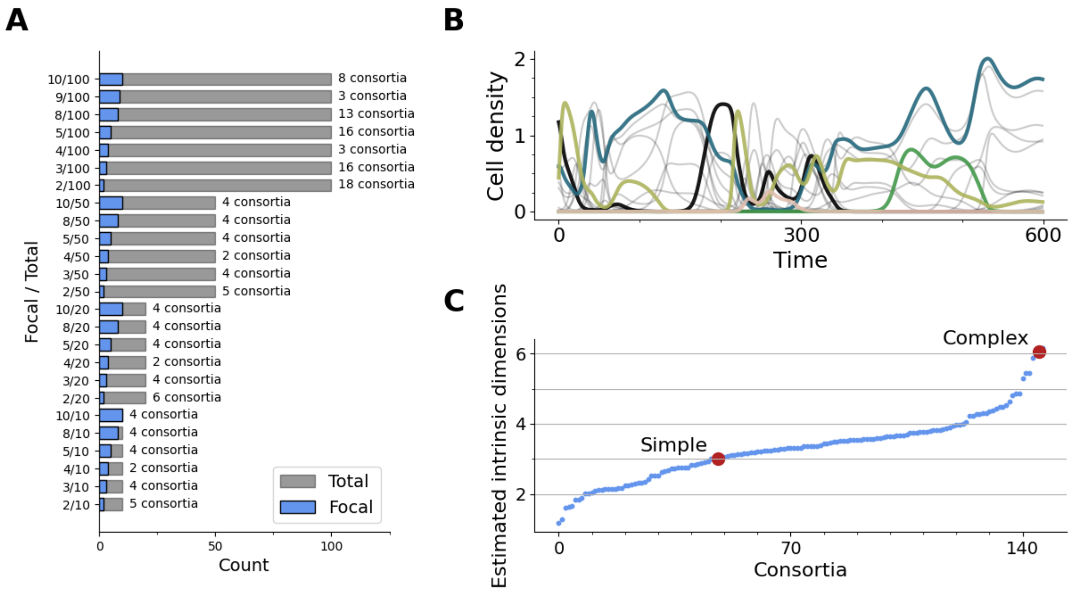


**Supplemental Figure S9. Description of the 150 simulated consortia**

A) The consortia for forecasting were simulated using three different consortia models. For each simulation, there was a total number of members in the simulation, and then a certain number of focal populations that were selected for analysis. There were 150 different consortia used in this analysis. Each consortium had a total number of members of 10, 20, 50, or 100. Each consortium had a focal community size of 2, 3, 4, 5, 8, or 10. For each of the unique consortia, there were 10,000 simulations. During training, 2,000 curves were used as the test set, and either 8,000 or 200 curves were used for the training set, depending on whether the model was a full or sparse dataset.

B) When modeling large consortia, we can focus on the populations of interest, and treat all the other strains as background. This is a sample community where the curves with color are the focal community and the gray curves are the background members.

C) Estimated intrinsic dimensions for all of the consortia, sorted in order of estimated intrinsic dimensions. We selected a consortium with estimated intrinsic dimensions of 3 as our simple community and 6 as our complex community.

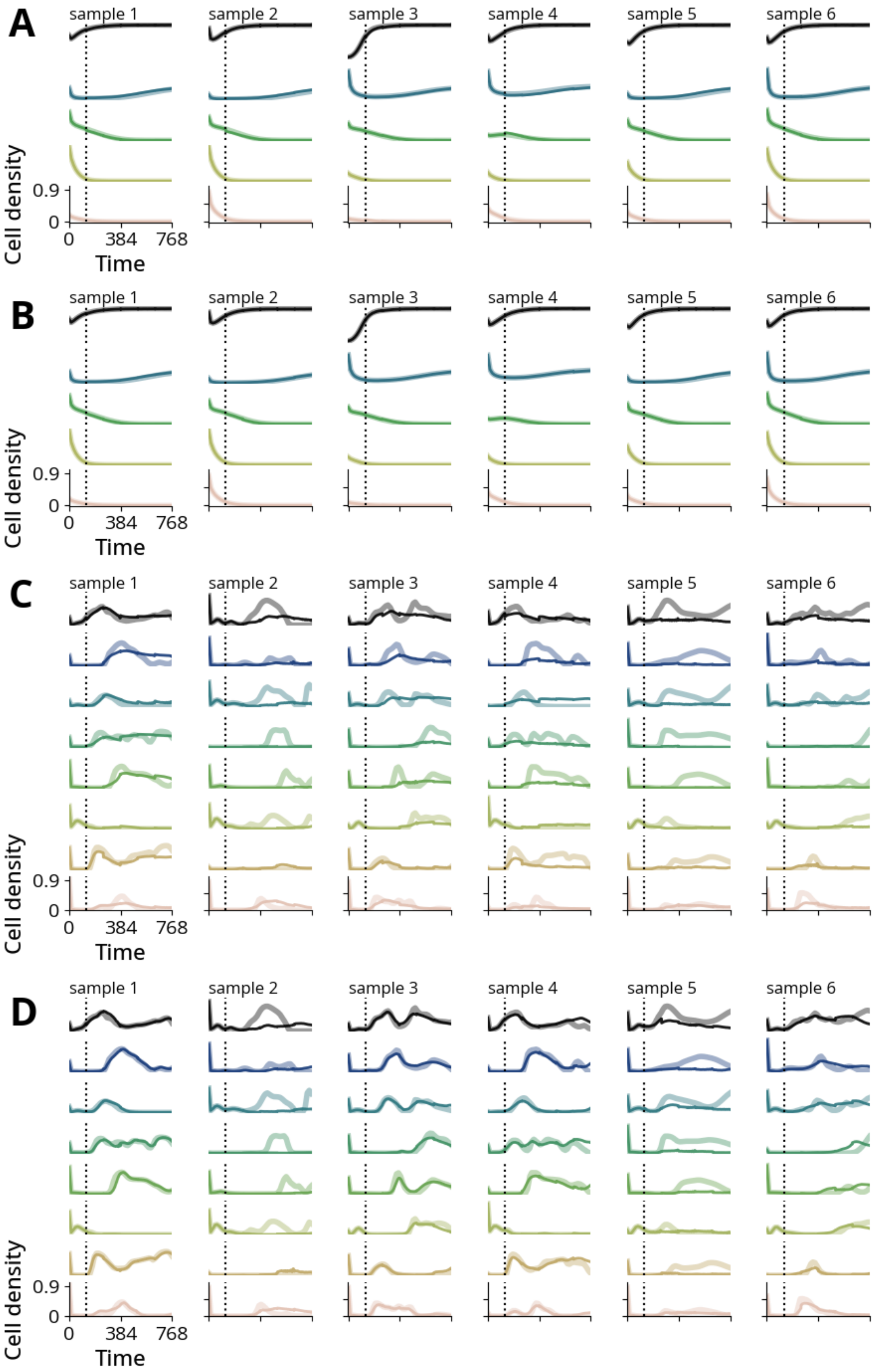
A
B
C
D
sample 1
sample 2
sample 3
sample 4
sample 5
sample 6
Cell density
0.9
0
0
384
768
Time

**Supplemental Figure S10. Sample forecast curves of the simulated consortia**

These curves are more samples of the simulated consortia for the main text. The models to predict the next segment were trained on either a full training dataset of 8,000 curves, or they were trained on a sparse training dataset of 200 curves. These results show using the latent representation of 128 time points to iteratively predict the next 128 time points.

A) Simple consortia trained on sparse training dataset. Samples are also used in B.
B) Simple consortia trained on full training dataset. Samples are also used in A.
C) Complex consortia trained on sparse training dataset. Samples are also used in D.
D) Complex consortia trained on full training dataset. Samples are also used in C.

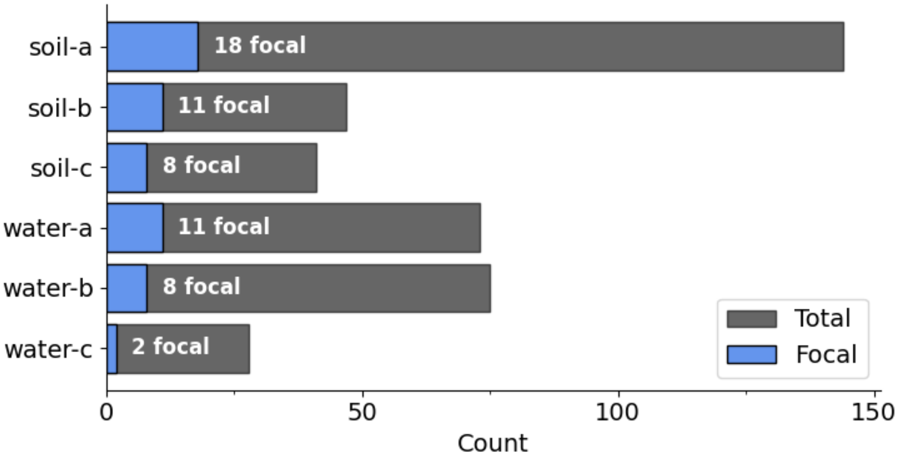


**Supplemental Figure S11. Summary of focal community sizes for the experimental consortia**

The experimental consortia dataset consisted of 6 experimental conditions with 8 replicates each. For each of the experimental conditions, there were 28 to 144 total members present across the 8 replicates. However, for each of the experimental conditions, only a subset of those members were present in all 8 replicates. These subsets are what we use as our focal communities, and their number of members ranged from 2 to 18.

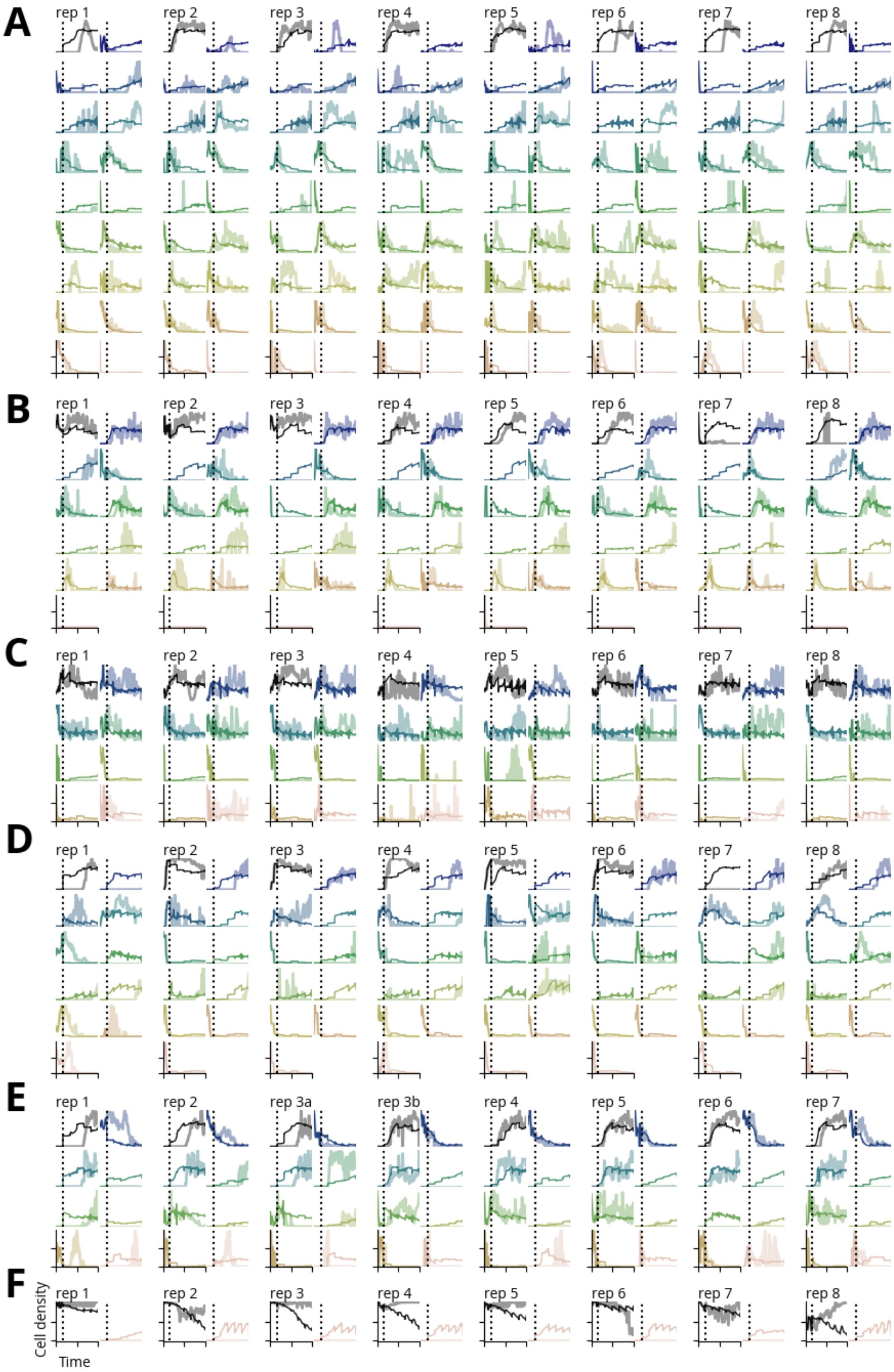
A
rep 1
rep 2
rep 3
rep 4
rep 5
rep 6
rep 7
rep 8
B
rep 1
rep 2
rep 3
rep 4
rep 5
rep 6
rep 7
rep 8
C
rep 1
rep 2
rep 3
rep 4
rep 5
rep 6
rep 7
rep 8
D
rep 1
rep 2
rep 3
rep 4
rep 5
rep 6
rep 7
rep 8
E
rep 1
rep 2
rep 3a
rep 3b
rep 4
rep 5
rep 6
rep 7
F
rep 1
rep 2
rep 3
rep 4
rep 5
rep 6
rep 7
rep 8
Cell density
Time

**Supplemental Figure S12. Sample forecasts for all experimental consortia**

These curves show the forecasted consortia for all of the 48 experiments in the dataset. For each replicate, the other 7 replicates were used to train the model to forecast the consortium. These forecasts were completed by encoding the initial 128 points into their latent representation, then predicting the latent representation for the following 128 points. Then, forecasts were made iteratively by appending each prediction to the initial segment.

A) Forecasts from the soil microbiome grown in oatmeal.
B) Forecasts from the soil microbiome grown in oatmeal-peptone.
C) Forecasts from the soil microbiome grown in peptone.
D) Forecasts from the water microbiome grown in oatmeal.
E) Forecasts from the water microbiome grown in oatmeal-peptone.
F) Forecasts from the water microbiome grown in peptone.

**Supplementary Table ST1. Simulation parameters for modified logistic equation**

We ran simulations using ordinary differential equations to model the modified logistic equation, which has been used in our previous studies [7], [8]. We ran 10,000 individual simulations for a duration of 24 (unitless time), with 128 time points per simulation. The parameters were randomly selected for each simulation using the ranges and distributions specified below. The ordinary differential equations were solved using the explicit Runge-Kutta method of order 5(4) via the scipy.integrate.solve_ivp function [9].

| Parameter | Description | Min | Max | Distribution |
|---|---|---|---|---|
| $n_0$ | Initial cell density | 0.03 | 0.07 | Uniform |
| $\alpha$ | Growth rate | 0.1 | 3.0 | Gamma |
| $n_{max}$ | Carrying capacity | 2.8 | 3.0 | Uniform |
| $k$ | Modification constant | 0.1 | 0.4 | Gamma |
| $\theta$ | Modification constant | 0 | 5 | Uniform |
| $t_{lag}$ | Lag time | 0 | 2 | Gamma |

**Supplementary Table ST2. Simulation parameters for chaotic consortia**

We ran simulations using ordinary differential equations to model the chaotic consortia equation. We ran 10,000 individual simulations for a duration of 500 (unitless time), with 128 time points per simulation. We repeated this 4 times. The parameters were randomly selected for each simulation using the ranges and distributions specified below. The ordinary differential equations were solved using the explicit Runge-Kutta method of order 5(4) via the scipy.integrate.solve_ivp function [9].

| Parameter | Description | Min | Max | Value | Distribution |
|---|---|---|---|---|---|
| *n0* | Initial cell density | n/a | n/a | 0.1 | Constant |
| $a_{i,j}$ | Interaction of species *i* with species *j* | -1 | 1 | n/a | Normal $N(0.3,0.25)$ |
| $a_{i,i}$ | Carrying capacity (interaction with self) | n/a | n/a | 1 | Constant |
| *M* | Number of members | n/a | n/a | 50 | Constant |
| *D* | Dispersal rate | n/a | n/a | 1e-6 | Constant |

**Supplementary Table ST3. Simulation parameters for β-lactam resistant**

We ran simulations using ordinary differential equations to model the β-lactam resistant equation. We ran 9,000 individual simulations for a duration of 16 (unitless time), with 128 time points per simulation. The parameters were randomly selected for each simulation using the ranges and distributions specified below. The ordinary differential equations were solved using the explicit Runge-Kutta method of order 5(4) via the scipy.integrate.solve_ivp function [9].

| Parameter | Description | Min | Max | Value | Distribution |
|---|---|---|---|---|---|
| $\alpha$ | Costs of β-lactamase production | 0.75 | 1.00 | n/a | Uniform |
| $\beta_{min}$ | Minimum benefit of β-lactamase production | 0.1 | 1.0 | n/a | Uniform |
| $\xi$ | Nutrient recycling | 0.1 | 1.0 | n/a | Uniform |
| $\kappa_b$ | Antibiotic degradation by β-lactamase | 0.1 | 1.0 | n/a | Uniform |
| $d_b$ | Effect of β-lactamase inhibitor on β-lactamase | 1 | 10 | n/a | Uniform |
| $\gamma$ | Lysis rate by antibiotic | 1.1 | 1.4 | n/a | Uniform |
| $i$ | Inhibitor concentration | 0.1 | 10 | n/a | Uniform |
| $\varphi_{max}$ | Maximum antibiotic degradation by living cells | 0.1 | 5 | n/a | Uniform |
| $c$ | Private benefit | 0.1 | 0.8 | n/a | Uniform |
| $n_{r,0}$ | Initial resistant cell density | 0.1 | 0.4 | n/a | Uniform |

| | | | | | |
|---|---|---|---|---|---|
| $n_{s,0}$ | Initial sensitive cell density | 0.1 | 0.4 | n/a | Uniform |
| $s_0$ | Initial nutrient concentration | n/a | n/a | 4 | Constant |
| $a_0$ | Initial antibiotic concentration | 1 | 100 | n/a | Uniform |
| $b_0$ | Initial β-lactamase concentration | n/a | n/a | 0 | Constant |
| $d_a$ | Basal antibiotic degradation | n/a | n/a | 0.02 | Constant |
| $h_i$ | Hill coefficient for inhibitor | n/a | n/a | 2 | Constant |
| $h_a$ | Hill coefficient for antibiotic | n/a | n/a | 3 | Constant |

**Supplementary Table ST4. Simulation parameters for bounded generalized Lotka-Volterra**

We ran simulations using ordinary differential equations to model the bounded generalized Lotka-Volterra equation. The parameters were randomly selected for each simulation using the ranges and distributions specified below.

| Parameter | Description | Min | Max | Distribution |
|---|---|---|---|---|
| *n0* | Initial cell density | 0 | 1 | Uniform |
| $\mu$ | Growth rate | 0 | 1 | Uniform |
| $\sigma$ | Environmental stress | 0.05 | 0.25 | Uniform |
| $\gamma^{+}$ | Positive interaction matrix | 0 | 1 | Uniform |
| $\gamma^{-}$ | Negative interaction matrix | 0 | 1 | Uniform |

**Supplementary Table ST5. Model comparison on reconstruction task**

We trained various models to accomplish the encoding task, and we are able to compare their performance on reconstruction accuracy. **Autoencoder 7X (A7X)**: variational autoencoder with 7 encoding layers and 8 decoding layers using a combination of linear and convolution layers; **Multi-layer perceptron Network Model (MNM)**: uses solely linear layers to build the variational autoencoder; **Principal components analysis Reducer (PR)**: uses traditional principal components analysis to reconstruct the curves; **MicroBERT Curve Reducer (MCR)**: based on the BERT transformer model; **Variational autoencoder Bottlenecked (VB)**: variational autoencoder bottleneck relying solely on convolution layers. All results presented are from the test datasets containing experimental data (exp), simulated data (sim), or both.

| **Model type** | **Test dataset** | **Reconstruction accuracy ($R^2$) by latent dimension** | | | | | | | | |
|---|---|---|---|---|---|---|---|---|---|---|
| | | **2** | **4** | **6** | **8** | **12** | **16** | **20** | **24** | **32** |
| A7X | both | 0.943 | 0.97 | 0.982 | 0.988 | 0.989 | 0.989 | 0.988 | 0.988 | 0.988 |
| A7X | exp | 0.946 | 0.969 | 0.981 | 0.987 | 0.987 | 0.987 | 0.987 | 0.987 | 0.987 |
| A7X | sim | 0.939 | 0.975 | 0.988 | 0.993 | 0.993 | 0.993 | 0.993 | 0.993 | 0.993 |
| MCR | both | 0.042 | 0.684 | 0.861 | 0.889 | 0.932 | 0.962 | 0.973 | 0.983 | 0.991 |
| MCR | exp | 0.042 | 0.665 | 0.858 | 0.882 | 0.925 | 0.959 | 0.973 | 0.983 | 0.991 |
| MCR | sim | -0.038 | 0.723 | 0.86 | 0.907 | 0.949 | 0.968 | 0.971 | 0.981 | 0.991 |
| MNM | both | 0.912 | 0.962 | 0.976 | 0.984 | 0.993 | 0.995 | 0.998 | 0.998 | 0.998 |
| MNM | exp | 0.914 | 0.96 | 0.974 | 0.982 | 0.991 | 0.995 | 0.998 | 0.998 | 0.998 |
| MNM | sim | 0.898 | 0.966 | 0.984 | 0.991 | 0.996 | 0.997 | 0.998 | 0.998 | 0.999 |

| | | | | | | | | | | |
|---|---|---|---|---|---|---|---|---|---|---|
| PR | both | 0.734 | 0.909 | 0.94 | 0.956 | 0.977 | 0.993 | 0.999 | 0.999 | 1 |
| PR | exp | 0.72 | 0.911 | 0.936 | 0.951 | 0.974 | 0.992 | 0.999 | 0.999 | 0.999 |
| PR | sim | 0.755 | 0.897 | 0.947 | 0.969 | 0.988 | 0.996 | 0.999 | 0.999 | 1 |
| VB | both | 0.944 | 0.975 | | 0.99 | | 0.997 | | | 0.999 |
| VB | exp | 0.947 | 0.975 | | 0.989 | | 0.997 | | | 0.999 |
| VB | sim | 0.941 | 0.976 | | 0.993 | | 0.999 | | | 1 |

**Supplementary Table ST6. Model comparison on classifying antibiotic type.**

We used all the models to classify the type of antibiotic used in the experiments presented in Fig 3A. The classification accuracy using the raw curves was 0.877. All data presented in this table are the mean accuracy of the test dataset from each of the 5 cross-validation folds.

| Model type | Classification accuracy by latent dimension | | | | | | | | |
|---|---|---|---|---|---|---|---|---|---|
| | 2 | 4 | 6 | 8 | 12 | 16 | 20 | 24 | 32 |
| A7X | 0.736 | 0.849 | 0.879 | 0.888 | 0.887 | 0.883 | 0.889 | 0.889 | 0.892 |
| MCR | 0.357 | 0.677 | 0.803 | 0.83 | 0.843 | 0.856 | 0.85 | 0.864 | 0.874 |
| MNM | 0.73 | 0.861 | 0.885 | 0.882 | 0.893 | 0.894 | 0.9 | 0.897 | 0.895 |
| PR | 0.723 | 0.847 | 0.884 | 0.89 | 0.886 | 0.889 | 0.888 | 0.89 | 0.886 |
| VB | 0.719 | 0.837 | | 0.887 | | 0.893 | | | 0.892 |

**Supplementary Table ST7. Model comparison on predicting antibiotic concentration.** We used all the models to predict the antibiotic concentration used in the experiments presented in Fig 3C. The prediction accuracy using the raw curves was 0.828. All data presented in this table are the mean accuracy of the test dataset from each of the 5 cross-validation folds.

| Model type | Prediction accuracy ($R^2$) by latent dimension | | | | | | | | |
|---|---|---|---|---|---|---|---|---|---|
| | **2** | **4** | **6** | **8** | **12** | **16** | **20** | **24** | **32** |
| A7X | 0.7 | 0.778 | 0.814 | 0.829 | 0.83 | 0.837 | 0.835 | 0.834 | 0.841 |
| MCR | -0.455 | 0.656 | 0.755 | 0.768 | 0.786 | 0.789 | 0.787 | 0.813 | 0.828 |
| MNM | 0.681 | 0.805 | 0.826 | 0.841 | 0.852 | 0.855 | 0.857 | 0.854 | 0.852 |
| PR | 0.697 | 0.789 | 0.822 | 0.841 | 0.851 | 0.859 | 0.859 | 0.855 | 0.855 |
| VB | 0.711 | 0.784 | | 0.802 | | 0.837 | | | 0.847 |

**Supplementary Table ST8. Model comparison on classification of resistance to CIP.**

We used all the models to classify the resistance of strains to CIP as presented in Fig 3E. The classification accuracy using the raw curves was 0.879. All data presented in this table are the mean accuracy of the test dataset from each of the 5 cross-validation folds.

| **Model type** | **Latent dimensions** | | | | | | | | |
|---|---|---|---|---|---|---|---|---|---|
| | **2** | **4** | **6** | **8** | **12** | **16** | **20** | **24** | **32** |
| A7X | 0.783 | 0.87 | 0.881 | 0.9 | 0.899 | 0.879 | 0.885 | 0.885 | 0.883 |
| MCR | 0.631 | 0.777 | 0.793 | 0.789 | 0.83 | 0.84 | 0.858 | 0.844 | 0.858 |
| MNM | 0.789 | 0.876 | 0.873 | 0.88 | 0.888 | 0.899 | 0.897 | 0.894 | 0.896 |
| PR | 0.79 | 0.837 | 0.858 | 0.875 | 0.891 | 0.897 | 0.9 | 0.896 | 0.887 |
| VB | 0.78 | 0.847 | | 0.893 | | 0.906 | | | 0.895 |

**Supplementary Table ST9. Model comparison on classification of resistance to GM.**

We used all the models to classify the resistance of strains to GM as presented in Fig 3E. The classification accuracy using the raw curves was 0.639. All data presented in this table are the mean accuracy of the test dataset from each of the 5 cross-validation folds.

| Model type | Latent dimensions | | | | | | | | |
|---|---|---|---|---|---|---|---|---|---|
| | 2 | 4 | 6 | 8 | 12 | 16 | 20 | 24 | 32 |
| A7X | 0.354 | 0.537 | 0.609 | 0.669 | 0.643 | 0.672 | 0.667 | 0.632 | 0.669 |
| MCR | 0.185 | 0.41 | 0.462 | 0.448 | 0.51 | 0.55 | 0.561 | 0.53 | 0.632 |
| MNM | 0.412 | 0.57 | 0.588 | 0.614 | 0.639 | 0.643 | 0.688 | 0.69 | 0.645 |
| PR | 0.351 | 0.551 | 0.568 | 0.627 | 0.636 | 0.613 | 0.558 | 0.569 | 0.547 |
| VB | 0.331 | 0.586 | | 0.648 | | 0.636 | | | 0.621 |

**Supplementary Table ST10. Model comparison on classification of resistance to SAM.**

We used all the models to classify the resistance of strains to SAM as presented in Fig 3E. The classification accuracy using the raw curves was 0.879. All data presented in this table are the mean accuracy of the test dataset from each of the 5 cross-validation folds.

| Model type | Latent dimensions | | | | | | | | |
|---|---|---|---|---|---|---|---|---|---|
| | 2 | 4 | 6 | 8 | 12 | 16 | 20 | 24 | 32 |
| A7X | 0.792 | 0.87 | 0.894 | 0.907 | 0.9 | 0.901 | 0.9 | 0.898 | 0.898 |
| MCR | 0.594 | 0.784 | 0.795 | 0.807 | 0.827 | 0.862 | 0.867 | 0.86 | 0.87 |
| MNM | 0.797 | 0.871 | 0.885 | 0.897 | 0.903 | 0.901 | 0.905 | 0.905 | 0.9 |
| PR | 0.804 | 0.853 | 0.883 | 0.892 | 0.895 | 0.901 | 0.891 | 0.892 | 0.893 |
| VB | 0.792 | 0.868 | | 0.906 | | 0.914 | | | 0.913 |

**Supplementary Table ST11. Model comparison on classification of resistance to SXT.**

We used all the models to classify the resistance of strains to SXT as presented in Fig 3E. The classification accuracy using the raw curves was 0.858. All data presented in this table are the mean accuracy of the test dataset from each of the 5 cross-validation folds.

| **Model type** | **Latent dimensions** | | | | | | | | |
|---|---|---|---|---|---|---|---|---|---|
| | **2** | **4** | **6** | **8** | **12** | **16** | **20** | **24** | **32** |
| A7X | 0.752 | 0.859 | 0.861 | 0.878 | 0.859 | 0.857 | 0.861 | 0.859 | 0.858 |
| MCR | 0.538 | 0.757 | 0.76 | 0.767 | 0.804 | 0.809 | 0.83 | 0.82 | 0.83 |
| MNM | 0.764 | 0.847 | 0.856 | 0.862 | 0.863 | 0.861 | 0.866 | 0.864 | 0.855 |
| PR | 0.768 | 0.825 | 0.845 | 0.857 | 0.862 | 0.86 | 0.868 | 0.863 | 0.86 |
| VB | 0.761 | 0.855 | | 0.874 | | 0.876 | | | 0.865 |

**Supplementary Table ST12. Experimental data**

We collected a variety of experimental datasets for this study. Many of these datasets were published and represent a broad array of different growth conditions and experiments. Additional datasets were gathered from our lab and others at Duke University. All of these datasets were completed following similar protocols to the previously published datasets, and the description of the methods for each of these can be found below.

| Source | Publication | Source | Publication |
|---|---|---|---|
| David *et al.* | [10] | Davis, H. | This study |
| Fujita *et al.* | [11] | Duncker, K.E. | This study |
| Ye *et al.* | [12] | Ha, Y. | This study |
| Hennigan *et al.* | [13] | Hamrick, G.S. | This study |
| Li *et al.* | [14] | Hoffman, A.L. | This study |
| Li *et al.* | [15] | Holmes, Z.A. | This study |
| Menacho Melgar *et al.* | [16] | Kim, K. | This study |
| Moreb *et al.* | [17] | Lee, D. | This study |
| Chory *et al*. | [18] | Liu, S. | This study |
| Schluter *et al.* | [19] | Lu, J. | This study |
| Maddamsetti, R | [20] | Shapiro, D.M. | This study |
| Ma *et al.*, Zhang *et al.*, | [6], [7], [8], [21] | Shende, A.R. | This study |

| | | | |
|---|---|---|---|
| Baig *et al.*, Ha *et al.* * | | | |
| Ahmad *et al.* | [22] | Villalobos, C.A. | This study |
| Aduru *et al.* | [23] | Wang, S. | This study |
| Palomino *et al.* | [24] | Yao, Z. | This study |
| Prensky *et al.* | [25] | | |
| Bethke *et al.* | [26] | | |
| Wu *et al*. | [27] | | |
| Şimşek, E. | [28], [29], and this study | | |

* Collection of growth curves run using Mantis robot and clinical isolates. The entire collection is referenced in Ma *et al*., and has been previously analyzed in other work in our group as seen in Zhang *et al.*, Baig *et al*., and Ha *et al.*

#### 2. Experimental methods

Folder: zach

**Bacterial strains and plasmids**

We used three collections of bacteria in our experiments. The clinical isolates were sourced from a collection of drug resistant bacteria from Duke University hospital and supplied by Vance Fowler and Joshua Thaden. Some of these strains have been further characterized in previous work in the lab [6], [26]. The bar-coded Keio strains were developed previously in our lab [27]. The sink isolates were collected from sinks at Duke University Hospital.

**Growth media and conditions**

The growth conditions for these experiments were completed as follows. Plates were made using LB agar. Cells were cultured in either 1) LB or 2) M9CA with 0.04% glucose. Overnight cultures were grown for ~20 to 24 hours in deep-well plates at 37°C and 700 RPM to allow the cells to reach the stationary phase. Cells were inoculated to reach exponential phase using a 16.67 μL to 1 mL ratio of overnight culture to fresh media, and then they were incubated for 2 hours in deep-well plates at 37°C and 700 RPM. We used kanamycin (Sigma) at 50 μg/mL to select for the plasmid-carrying Keio strains.

**Robotic liquid handling**

We used the Tecan Freedom EVO liquid handling robot to perform our experiments. The robot performed the dilutions from the exponential cultures to the 1,000× and 10,000× dilutions. Using the incubator in the robot, the cells were shaken and incubated at 37°C. The robot performed the plate reading by moving plates to the attached Tecan Infinite 200 PRO plate reader.

Folder: alex

**Bacterial strains and plasmids**

The bar-coded Keio strains were developed previously in our lab [27].

**Growth media and conditions**

The growth conditions for these experiments were completed as follows. Plates were made using LB agar. Cells were cultured in either 1) LB or 2) M9CA with 0.04% glucose. Overnight cultures were grown for ~20 to 24 hours in deep-well plates at 37°C and 700 RPM to allow the cells to reach the stationary phase. Cells were inoculated to reach exponential phase using a 16.67 μL to 1 mL ratio of overnight culture to fresh media, and then they were diluted and applied to different chemical combinations. We used kanamycin (Sigma) at 50 µg/mL to select for the plasmid-carrying Keio strains.

Folder: ashwini

**Bacterial strains and plasmids**

We used several strains in these experiments, all with an *Escherichia coli* MG1655 background. Strains contained the following plasmid combinations: either (1) FHR + ptet(RBS32)-Cas9-ssrA-SC101, or (2) FHR + ptet(RBS32)-Cas9-ssrA-SC101 + ptet-sp1-plac-sfGFP-ColE1-oriTF

**Growth media and conditions**

The growth conditions for these experiments were as follows: Overnight cultures were grown in either (1) LB + 25 ug/mL chloramphenicol or (2) LB + 25 ug/mL chloramphenicol + 50 ug/mL kanamycin. Overnights were grown in 3 mL culture tubes at 37°C and 700 RPM, for ~16.5 hours. Cultures were then diluted to an OD of 0.5. For some experiments, cultures from each strain were combined, while in others, single population (clonal) experiments were conducted. In all cases, 1 uL of diluted culture was inoculated in 1 mL LB + 25 ug/mL chloramphenicol for a final dilution rate of 1/1000. Various concentrations of anhydrous tetracycline (ranging from 0 ng/mL to 100 ng/mL) were added to cultures to induce plasmid cutting. Finally, each sample was distributed (200 µl per well) into a black-walled 96-well plate (Corning, CLS3603). Data was collected using a Tecan Infinite 200 PRO plate reader. The reader held plates at 37°C and took measurements of cell density (OD600) and GFP intensity (excitation: 488 nm, emission: 510 nm) every 10 minutes for 16-24 hours.

Folder: dongheon

**Bacterial strains and plasmids**

E. coli strain, DA28202 [30], was used for all experiments. Two plasmids (ptetNahRAM-mRuby3 and ptetNahRAM-mR3-IDP) were constructed from parent plasmids, p15aTetT7mut-fuseYFP [31], p-[WT]-20-mRuby3 [32], pAJM.771 [33] through Gibson assembly (NEBuilder HiFi DNA Assembly Master Mix). psalsfGFP were constructed from parent plasmids psfGFP, which was constructed by replacing pCFP [31] with sfGFP gene fragment synthesized by Integrated DNA Technologies. Subsequently, the T7 promoter in the psfGFP plasmid was replaced by psal promoter [33] with NEB Q5 Site-Directed Mutagenesis Kit.

**Measurements**

Cells carrying the plasmids were grown in 3mL of LB medium at 37°C at 225 RPM. After 16 hours of growth, the cell culture was diluted 100-fold in M9CA media supplemented with 0.4% glucose, 1mM thiamine hydrochloride, and appropriate antibiotics. Then, the cell culture was incubated at 37°C and 225 RPM. After 1.5 hours of incubation, the cell was distributed (200 µL per well) into a black-walled 96-well plate (Corning, CLS3603), and each well was added with varying amounts of aTc from 0 to 100 ng/mL. Additionally, 20 µL of mineral oil was added to each well. Cell density ($OD_{600}$), mRuby3 intensity (excitation: 561 nm, emission 560 nm), and GFP intensity (excitation: 470 nm, emission 515 nm) were measured using a microplate reader (Tecan Infinite 200 PRO). The plate was incubated at 37°C for 20 hours.

Folder: sizhe

**Bacterial strains and plasmids**

E. coli strain, MC4100Z1, containing ePop autolysis circuit [34] was used for all experiments. Plasmid p15A-t5-CEC was constructed from the parent plasmid, Cdc19-ELP35-Cdc19 (from Ashutosh Chilkoti's lab) through Gibson assembly (NEBuilder HiFi DNA Assembly MasterMix). Plasmid p15A-His-T3 was from lab previous work [35]. Two plasmids (p15A-t5-mCherry, p15A-t5-mCherry-Cdc19) were constructed from plasmid p15A-t5-CEC through Gibson assembly.

**Growth media and conditions**

The growth conditions for these experiments were completed as follows. Plates were made using LB agar supplemented with 2% glucose and appropriate antibiotics. 25 ug/mL chloramphenicol + 50 ug/mL kanamycin were used to select strains containing plasmids during the whole experiments. Cell colonies carrying the plasmids were grown in 3mL of LB medium supplemented with 2% (w/v) glucose at 37°C at 225 RPM. 1 mM IPTG and antibiotics were added to induce the target protein's expression. After ~18 hours of growth, the cell culture was diluted 100-fold in LB media, 1mM IPTG, and appropriate antibiotics. Different concentrations of glucose (0%, 0.2%, 2%) were added into media to control the ePop circuit. The cell was then distributed (200 µL per well) into a black-walled 96-well plate (Corning, CLS3603), and 50 µL of mineral oil was added to the top of each well. Data was collected using a Tecan Infinite 200 PRO plate reader. The reader held plates at 30/37/42°C (mostly 37°C) and took measurements of cell density (OD600) and mCherry intensity (excitation: 561 nm, emission: 600 nm) every 5 minutes for 16-28 hours.

Folder: cesar

*E. coli* of different β-lactam resistant or sensitive strains were cultured under varying concentrations of β-lactamase inhibitors - clavulanic acid, tazobactam, or sulbactam - without β-lactamase treatment. These were grown for 23 hours, at 37°C and moderate agitation in a Tecan Infinite 200 PRO, with an OD600 and fluorescence (if applicable) reading every 10 minutes. Sometimes Kanamycin was added to the outgrowth media, as indicated in the dataset. Some wells had no inhibitor and only carbenicillin treatment, to verify sensitivity to β-lactams.

For the experiments with resistant Top10 or DH5α, these were transformed with the β-lactamase-producing plasmid pBla [8].

Folder: zhixiang

**Bacterial Strains and Plasmids**

E. coli strains MG1655 and DA28102 were used for all experiments. Three plasmids, pUC, p15A, and ColEI, were included in the study. These plasmids represent various origins of replication and compatibility groups.

**Experimental Chemicals**

Rifampicin: 4 µg/mL

Linoleic acid: 40 µg/mL

Trovafloxacin: 0.032 µg/mL

Promethazine: 32 µg/mL

Maprotiline: 64 µg/mL

Phenothiazine: 16 µg/mL

Ascorbic Acid: 40 mM

Sodium Dodecyl Sulfate (SDS): 0.02%

**Culture Conditions**

Cells were grown in either M9CA media supplemented with 0.4% glucose or LB medium. For all conditions, a control group with carbenicillin at 100 µg/mL was included.

**Growth and Testing Procedure**

Cultures of E. coli MG1655 and DA28102 strains carrying pUC, p15A, and ColEI plasmids were started in M9CA medium supplemented with 0.4% glucose or LB medium. The initial culture was grown for 16 hours at 37°C with shaking at 225 RPM. Before incubation in black-walled 96-well plates, cells were first incubated in a tube with 5 mL media containing carbenicillin (100 µg/mL), shaking at 225 RPM, at 37°C. For experiments with DA28102, 25 µg/mL chloramphenicol was also added to all groups. Carbenicillin was used to select for plasmids, and chloramphenicol was used to select for DA28102 strains. Cultures were diluted daily at a rate of 1:200 for 3-15 days, with daily measurements taken at 20-23.5-hour intervals. Chemical conditions tested included single chemical groups and double chemical combinations, as specified above. Cultures were transferred into black-walled 96-well plates (Corning, CLS3603) for measurements, with each well containing 200 µL of the respective culture. For all wells, 20 µL of mineral oil was added to prevent evaporation.

**Measurements**

Cell density and GFP fluorescence were measured using a Tecan Infinite 200 Pro plate reader at 30-minute intervals over varying total lengths of time (ranging from 20 to 23.5 hours): Cell density: OD600, GFP fluorescence: Excitation at 470 nm, emission at 515 nm.

**Testing Equipment**

All measurements were conducted using a Tecan Infinite 200 Pro plate reader. Plates were incubated at 37°C during the measurement periods.

**Control Conditions**

Control groups included cultures treated with carbenicillin at 100 µg/mL to ensure baseline comparisons.

Folder: dan

**Protocol 1**

Grew up in 2xYT + Kanamycin + chloramphenicol from frozen stock overnight, then back-diluted 1uL into 180uL M9CA +Kan +Cat +1% glucose + 1mM IPTG or .2% arabinose and recorded in plate reader for the following curves. Used for the mCherry part of file “22_2_7 mCherry and Protein E vs time with Pum2-ELP-GFP constructs.csv”.

**Protocol 2**

Grew up in 2xYT + Kanamycin + chloramphenicol from frozen stock overnight, back-diluted 50uL into 2mL (OD600 0.1 typically), grew for 3 hours, then recorded 180uL in plate reader with or without induction. Used for the protein E part of file “22_2_7 mCherry and Protein E vs time with Pum2-ELP-GFP constructs.csv”.

**Protocol 3**

Grew up in MOPS EZ rich medium [36] overnight from frozen stock, then back-diluted 100uL into 2mL (OD600 0.2), grew for 3 hours, then recorded 180uL in plate reader with or without induction.

**Protocol 4**

Grew up overnight in 2xYT + Kan + Chloramphenicol + 1% glucose overnight, BD to OD600 0.1, grew until 0.3, added arabinose, grew until 0.8, switched to IPTG, then put into plate reader.

Folder: katie

**Sensor strains, plasmids, and experiments**

<u>Sensor strains and plasmids</u>

The IPTG and aTc sensor pair were composed of p15a-T5-2A-mCherry and p15a-plTetO1-sfGFP plasmids, respectively, which were both kanamycin resistant and were contained in *E. coli* MC4100Z1 host strains. p15a-T5-2A-mCherry plasmid was obtained from a previous study, in which it was called His-$T_2$-mCherry [35]. p15a-pTet-sfGFP was modified from the p15A-pTet-sfGFP-linker-Tdimer-kanR, from which the linker-Tdimer sequence was removed [37].

The thiosulfate (THS) and tetrathionate (TTR) sensor circuits were using the following plasmids pKD236-4b, pKD237-3a-2, pKD238-1a, and pKD239-1g-2 which were gifts from Jeffrey Tabor [38] (Addgene plasmid # 90956 ; http://n2t.net/addgene:90956 ; RRID:Addgene_90956), (Addgene plasmid # 90957 ; http://n2t.net/addgene:90957 ; RRID:Addgene_90957), (Addgene plasmid # 90958 ; http://n2t.net/addgene:90958 ; RRID:Addgene_90958), (Addgene plasmid # 90959 ; http://n2t.net/addgene:90959 ; RRID:Addgene_90959). The plasmids containing the regulator gene and fluorescence reporter gene (pKD237-3a-2 and pKD239-1g-2) were modified to have a different fluorescent reporter (CFP and YFP, respectively) in place of sfGFP, and the constitutively expressed mCherry was removed. The mCherry, YFP, and CFP gene sequences were obtained from plasmids previously developed in our lab. The other plasmids were unmodified. The two plasmids corresponding to each two-component system sensor were co-transformed into *E. coli* MG1655 strain via CaCl2 transformation and plated on LB agar plates with chloramphenicol and spectinomycin selection.

The N-(3-Hydroxytetradecanoyl)-DL-homoserine lactone (OHC14), cuminic acid (Cuma), and aTc sensors were constructed from the following plasmids pAJM.1642, pAJM.657, and pAJM.011, which were a gift from Christopher Voigt [33] (Addgene plasmid # 108535 ; http://n2t.net/addgene:108535 ; RRID:Addgene_108535), (Addgene plasmid # 108525 ; http://n2t.net/addgene:108525 ; RRID:Addgene_108525), (Addgene plasmid # 108529 ; http://n2t.net/addgene:108529 ; RRID:Addgene_108529). The YFP reporter was replaced with mCherry in the aTc sensor (pAJM.011) and with CFP in the OHC14 sensor (pAJM.1642). The plasmids were transformed into *E. coli* Top10F' background strain via CaCl2 transformation and plated on LB agar plates with kanamycin selection.

Plasmid modifications were performed using polymerase chain reaction (PCR) and Gibson Assembly cloning method [39]. Colonies from the transformation were cultured overnight in LB with corresponding antibiotic selection and stored in 25% glycerol at -80°C.

<u>Sensor experiment protocol:</u>

Freezer stocks were streaked onto LB agar plates with appropriate antibiotic selection and incubated at 37°C overnight. Single colonies were picked and cultured overnight in 3mL LB media with appropriate antibiotics. OD was measured and cultures were diluted to reach the same OD. The two or three sensor strain cultures were either combined at an equal ratio and distributed to 96-well plates (Corning, black-sided, clear flat bottom) or individually distributed to 96-well plates for co-culture and mono-culture experiments, respectively. Corresponding antibiotics and inducers were added at various concentrations to the 96-well cultures. The inducers used consist of the following: IPTG, aTc (Sigma), sodium thiosulfate pentahydrate (Sigma), potassium tetrathionate (Sigma), N-(3-Hydroxytetradecanoyl)-DL-homoserine lactone (Sigma), 4-Isopropylbenzoic acid (Cuma) (Sigma). Plates were incubated overnight with periodic plate readings in Tecan Infinite 200 PRO. Some experiments used the Tecan Freedom EVO robot for moving multiple plates from incubator to plate reader periodically.

**Growth rate strains, plasmids, and experiments:**

<u>Sink strains</u>

Sink strains were obtained from Helena Ma (box 2 and box 3). Water was collected from hospital sink p-traps and either directly plated on LB agar plates overnight or cultured in liquid culture before culturing on LB agar plates. Colonies were picked, cultured overnight, and then stored at -80°C. For the experiments in this study, individual strains were streaked from freezer stocks onto LB agar plates and incubated at 37°C overnight. Individual colonies were picked and inoculated into tryptic soy broth for overnight culturing until cultures reached stationary phase. Stationary phase cultures were diluted 1:60 in tryptic soy broth and cultured for 3 hours to reinstate exponential phase. Exponential phase cultures were diluted 1,000-fold, 10,000-fold, and 100,000-fold and distributed to a 96-well plate. The wells were covered with 50uL mineral oil to prevent evaporation and incubated overnight with periodic plate readings in Tecan Infinite 200 PRO plate reader.

Amyloid strains

All strains used in this experiment were *E. coli* MC4100Z1 host strains containing ePop autolysis circuit [34] as well as one of the following plasmids: p15A-t5-CEC, p15A- His-T3 [35], and p15A-t5-mCherry-Cdc19. Plasmid p15A-t5-CEC was constructed from the parent plasmid, Cdc19-ELP35-Cdc19 (in collaboration with Max Ney and Ashutosh Chilkoti), and p15A-t5-mCherry-Cdc19 was modified from p15A-T5-CEC. All plasmid modifications were done through Gibson assembly (NEBuilder HiFi DNA Assembly MasterMix) [39]. The experimental protocol is as follows: strains were streaked on LB agar plates with 2% glucose, chloramphenicol and kanamycin; colonies were cultured overnight in 3 mL LB with 2% glucose, antibiotics, and IPTG at 37°C shaking conditions; cultures were diluted 1,000-fold and 10,000-fold and distributed to a 96-well plate for monocultures and co-cultures. Conditions varied in this experiment include media pH at 4.8, 5.8, or 7 and glucose concentration at 0% or 0.2% w/v. 200uL cultures were covered with 50uL mineral oil to prevent evaporation and incubated overnight with periodic plate readings in Tecan Infinite 200 PRO plate reader.

Folder: grayson and hyein

**96-well plate growth curves**

Cells were grown overnight (16 hours, 37°C, 700 RPM, 1 mL/well) in LB media without antibiotics in a 2 mL deep 96-well plate. Cultures were diluted 1,000-fold in fresh LB broth and transferred into a flat-bottom 96-well plate (200 μl per well final volume, VWR). To prevent evaporation, 50 μl of mineral oil (Sigma Aldrich, SKU M5904) was added to the top of each well. Timelapse optical density (600 nm) was measured every 10 minutes for at least 12 hours using a plate reader (Tecan Infinite 200 PRO). The plate was shaken in an orbital motion (radius = 2 mm) for 5 seconds before each reading.

**1536-well plate growth curves**

Cells were grown overnight (16 hours, 700 rpm, 1 mL/well) in LB at 37°C (E. coli) or Marine Broth at 30 ℃ (marine isolates) without antibiotics in a 2 mL deep 96-well plate. Cultures were diluted 1,000-fold in the appropriate fresh broth and transferred into a flat-bottom 1536-well plate (7 µl per well final volume, VWR) using a MANTIS automated liquid handler. Timelapse optical density (600 nm) was measured every 10 minutes for at least 12 hours using a plate reader (Tecan Spark). To prevent evaporation, the plate was loaded into the plate reader with a magnetic cover which was removed automatically for each measurement and was maintained at 30°C for both E. coli and marine isolates. The plate was shaken in an orbital motion (radius = 2 mm) for 5 seconds before each reading.

Folder: harris

**Bacterial strains**

Six strains were selected from the PA14 *Pseudomonas aeruginosa* non-essential transposon knockout library based on colony phenotype on solid swarming media: wildtype, plus MutantIDs 27109, 27146, 29423, 34276, 35588 [40], [41].

**Stock solution preparation**

Phosphate (PO4) buffer (5×) was prepared with 12.0 g dibasic sodium phosphate (MilliporeSigma, S3264), 15.0 g monobasic potassium phosphate (MilliporeSigma, P5655), and 2.5 g sodium chloride (MilliporeSigma, S3014) into 1 L distilled water, then autoclaved. All other solutions—casamino acids (ThermoFisher, 223120), magnesium sulfate (MilliporeSigma, M2643), calcium chloride (MilliporeSigma, C1016), and gentamicin (ThermoFisher, 15750060)—were prepared by dissolving appropriate mass in water and then sterile filtering.

**Growth media and conditions**

Glycerol stocks for each strain were kept at -80°C. Before each experiment, glycerol stocks were grown overnight in liquid LB (Genesee, 11-120) at 37°C and then streaked on solid LB/agar (Genesee, 11-122) to isolate colonies, which were grown overnight at 37°C. Two single colonies were picked as biological replicates and grown overnight in the same liquid LB at 37°C. Note that, as the PA14 mutants are gentamicin selectable, overnight cultures on liquid and solid media were grown with 15 µg/mL gentamicin for mutants. The wildtype strain was grown in the absence of gentamicin. Cultures for growth curves were prepared from the overnight liquid cultures of single colonies the following morning. Overnight cultures were diluted to 0.1 < OD600 < 0.2 and incubated for 1-2 hours. Liquid media was prepared with varying concentrations of casamino acids, $MgSO_4$, $CaCl_2$, $PO_4$ buffer, and gentamicin (gentamicin was omitted for wildtype cultures). Cultures were then centrifuged in 15 mL centrifuge tubes (Avantor/VWR, 21008) for 5 minutes at 3,000 rpm, 4°C. LB media was aspirated and cells were resuspended in 1 mL liquid media. Cells were then diluted to OD600 = 0.2. Cells were finally diluted with liquid media into an acrylic 96-well plate (Avantor/VWR, 29442) in 1:10 cells-to-total-volume to an assumed OD600 = 0.04. Water was added to perimeter wells of plate and a single blank well of just liquid media was also added. Wells containing any liquid volume were covered with 50 mL mineral oil (MilliporeSigma, M5904) and OD600 was read for 24 hours, shaken every 10 minutes, at 37°C.

Folder: yuanchi

**For tet data**

All experiments were conducted using E. coli strain MG1655. There are five different strains, each with a different plasmid, pSC101, p15A, CloDF13, pBR322 and pUC. For all experiments, we cultured cells in LB media (LB broth Miller mix from Apex Bioresearch Products) at 37°C at 225 RPM (for round-bottom culture tubes). For each plate, we either supplemented tetracycline selection or no antibiotic selection in all wells. For the plates with selections, we used one of the two tetracycline concentrations: 5 µg/mL or 10 µg/mL. We sealed the plate top with AeraSeal film sealant (Sigma-Aldrich) and measured the datapoint every 10 min after shaking in plate reader (Tecan Infinite 200 PRO) for 16 or 24 hours. Cell density (optical density at 600 nm wavelength, or OD600) and GFP intensity (Ex: 488 nm, Em: 510 nm) were measured for each timepoint.

**For inc data**

All experiments were conducted using E. coli strain MG1655. There are four different strains, each with a different plasmid, pSC101, p15A, ColE1 and pUC. For all experiments, we cultured cells in LB media (LB broth Miller mix from Apex Bioresearch Products) at 37°C at 225 RPM (for round-bottom culture tubes). For each plate, we either supplemented 25 µg/mL chloramphenicol selection or 50 µg/mL kanamycin antibiotic selection in all wells. We sealed the plate top with AeraSeal film sealant (Sigma-Aldrich) and measured the datapoint every 10 min after shaking in plate reader (Tecan Infinite 200 PRO) for 16 or 24 hours. Cell density (optical density at 600 nm wavelength, or OD600), GFP intensity (Ex: 488 nm, Em: 510 nm) and mCherry intensity (Ex: 561 nm, Em: 610 nm) were measured for each timepoint.

Folder: emrah

In every experiment, bacteria were first streaked on a Luria-Bertani (LB) agar plate from their frozen glycerol stock, and a single colony was incubated overnight in liquid LB media at 37°C and 225 r.p.m.

**Bacterial growth media and procedures for Group 1 and Group 2 data below:**

For all experiments, PBS (2.4 g/l $Na_2HPO_4$ (anhydrous), 3 g/l $KH_2PO_4$ (anhydrous), 0.5 g/l NaCl, 1 mM $MgSO_4$, and 0.1 mM $CaCl_2$) supplemented with casamino acids (8 g/l, unless otherwise noted) was used as the experimental media.

The media was prepared following a recipe adapted from Xavier *et al*. [42]: To make 1 liter of 5× phosphate buffer stock solution, 12 g $Na_2HPO_4$ (anhydrous), 15 g $KH_2PO_4$ (anhydrous), and 2.5 g NaCl were dissolved in deionized water and sterilized by autoclaving. The casamino acids stock solutions were made at 20 % (w/v) by microwaving 40 g casamino acids (Gibco™ Bacto™ 223120) in deionized water, sterilized by filtering (0.22 μm) and stored at 4°C. Then, appropriate amounts were immediately used to prepare the experimental media.

**Group 1:** Experiments dated 2022-11-30, 2023-05-22, 2023-05-30, and 2023-05-31

We used *P. aeruginosa* PA14 wild type, or a *fleN* mutant, or one of the swarming deficient variants identified as #36, #49, #58, #75, #87 in Luo, *et al*. [28].

We measured the growth curves of *P. aeruginosa* using a plate reader (Tecan Infinite 200). In each well of a 96-well plate, we added 200 μl liquid media and 2 μl cell culture (overnight cultures of different replicates and strains were diluted 10×~50× to the same cell density). To prevent evaporation, we added 50 μl mineral oil to each well. The cells were then incubated in the plate reader at 37°C, and $OD_{600}$ measurements were taken at 10-min intervals for 24 hours. Background signals were measured from media containing no cells (blank). For replicates of a particular strain, cells of the same stock were inoculated into different wells of a 96-well plate and measured.

The same data were also used in Luo, *et al*. [28].

**Group 2:** Experiments dated 2024-05-28, 2024-06-05, 2024-07-05, and 2024-07-11

2024-05-28 and 2024-06-05: *P. aeruginosa* PA14 wild type, or a clinical *K. pneumoniae* isolate D-005 from Deverick J. Anderson's laboratory, or a mixture with the 1:1 initial biomass ratio of the two was used. The bacteria were tested without or with cefotaxime (2.5 μg/ml treatment).

2024-07-05 and 2024-07-11: Hospital sink isolates *P. aeruginosa* 1926221, or *Bacillus paranthracis* 1926211 in Şimşek, *et al*., 2025, *bioRxiv* [29], or a mixture with the 1:1 initial biomass ratio of the two was used. The bacteria were tested without or with carbenicillin (10 μg/ml treatment).

We measured the growth curves of bacteria using a plate reader (Tecan Infinite 200). In each well of a 96-well plate, we added 195 μl liquid media and 5 μl cell inoculum culture (prepared by centrifuging overnight-grown cells at 1150 g for 5 min and then resuspending in the liquid medium). To prevent evaporation, we used Nunc Edge multi-well plates (Thermo Scientific) with built-in water reservoirs. The cells were then incubated in the plate reader at 37 °C, and $OD_{600}$ measurements were taken at 10 min intervals. For technical replicates of a particular condition, cells from the same inoculum were inoculated in different wells of a 96-well plate and measured. Background signals were measured from media containing no cells.

The same data were also used in in Şimşek, *et al*., 2025, *bioRxiv* [29].

**Group 3:** all others, which are unpublished.

We used *P. aeruginosa* PA14 wild type, or *E. coli* MC4100Z1 with a plasmid that encodes an engineered cytoplasmic beta-lactamase (BlaM:$Kan^R$) [43], or *E. coli* MC4100Z1 with the QS-CAT:$Kan^R$-$Cb^R$) circuit [44].

**Bacterial growth media and procedures for Group 3:**

*P. aeruginosa* PA14 wild type was tested in LB with varied concentrations of carbenicillin, or piperacillin tazobactam combinations, or piperacillin sulbactam combinations, or in 0.2xLB with varied concentrations of carbenicillin, at 30°C.

*P. aeruginosa* PA14 wild type was also tested in LB with varied concentrations of ticarcillin, or ceftazidime and clavulanic acid combinations, or in M9 with glucose (0.2 %) without or with casamino acids (0.2 %) together with varied concentrations of piperacillin, at 37°C.

*E. coli* MC4100Z1 BlaM was always cultured in LB with kanamycin (50 μg/ml) and IPTG (1 mM). Its growth was tested with varied concentrations of carbenicillin, at 37°C.

*E. coli* MC4100Z1 QS-CAT was always cultured with kanamycin (50 µg/ml) and carbenicillin (100 µg/ml). Its growth was tested with varied concentrations of chloramphenicol, at 37°C.

We measured the growth curves of bacteria using a plate reader (Tecan Infinite 200). In each well of a 96-well plate, we added 200 µl liquid media and 2 µl cell inoculum culture (prepared by centrifuging overnight-grown cells at 1150 g for 5 min and then resuspending in the liquid medium). To prevent evaporation, we added 50 µl mineral oil to each well. The cells were then incubated in the plate reader, and $OD_{600}$ measurements were taken at 10 min intervals. For technical replicates of a particular condition, cells from the same inoculum were inoculated in different wells of a 96-well plate and measured. Background signals were measured from media containing no cells.

Folder: kyeri

In every experiment, bacteria were first streaked on a Luria-Bertani (LB) agar plate from their frozen glycerol stock, and a single colony was incubated overnight in liquid media at 37°C and 225 RPM.

**KeioV3_mcherry files:**

The bar-coded Keio *E. coli* strains were developed previously in our lab [27], [45]. Cells were inoculated in media and grown in a plate reader. Readings were paused after an initial growth period to add antibiotics. Across the different experiments, there were 3 different antibiotics used: carbenicillin, amoxicillin, and cefotaxime. The antibiotics were applied at different concentrations as specified in the files.

**TOP10F files:**

The *E. coli* TOP10F cells were inoculated in different media and antibiotic concentrations with different starting cell densities. Cells were inoculated in media and grown in a plate reader.

**FJTimer_MG1655_cas_betalactams files:**

The *E. coli* MG1655 cells contained the p15A-pTet-sfGFP-linker-Tdimer-kanR plasmid [37], [46]. Cells were inoculated in media and grown in a plate reader. The media contained different casamino acid concentrations. Readings were paused after an initial growth period to add antibiotics. Across the different experiments, there were 3 different antibiotics used: carbenicillin, amoxicillin, and cefotaxime. The antibiotics were applied at different concentrations as specified in the files.

Folder: zach_other

**Bacterial strains**

In this work the bacterial strains used included *E. coli* strains MC4100Z1, TOP10F', MG1655, DH5α.

**Media, antibiotics, and chemicals**

The cells were cultured in lysogeny broth (LB) media. The antibiotics used were chloramphenicol (34 μg mL$^{-1}$) and kanamycin (50 μg mL$^{-1}$). Glucose was used at 0.2% concentration. Isopropyl β-d-1-thiogalactopyranoside (IPTG) was added at 1 mM concentration. Other chemicals were added as specified in specific files.

**Plasmids**

The ePop circuit was published in previous work from the group [34]. T5-p15A-GFP-OLS and T5-p15A-mCherry-OAC were constructed by replacing genes in T5-p15A-pAAA from previous work in the group [35]. The OLS and OAC genes were ordered from Integrated DNA Technologies. The GFP was cloned from a plasmid from our group. The mCherry was cloned from a plasmid T5-15A-mCherry-pAA from previous work in the group [35]. The BlaM, QS-CAT, and QS-BlaM were from previous work in the group [44].

**Culture**

Cells were cultured overnight in 2 mL or 3 mL of LB media at 37°C and shaking at 225 RPM.

**Evaluation of growth and fluorescent protein expression**

Generally, for long-term plate reader experiments, the cells were diluted 1000× into 1 mL of LB. The appropriate antibiotics, glucose, and chemicals were added. The cultures were then aliquoted into a 96-well plate in triplicates (200 µL per culture). To prevent evaporation, the cultures were covered with 50 µL of mineral oil. A plate reader (Tecan) was used to measure the optical density (OD600) and fluorescence signals (GFP and mCherry). Measurements were taken every 5 minutes for 40 to 48 hours. Measurements were corrected against the average of three blank samples in each run.

Folder: jia

**Bacterial strains**

There are 96 isolates by spatial evolution that all came from the same ancestry strain: *P. aeruginosa* PA14 wild type [28]. Each strain carries one or two types of mutations in their genome, and there are 18 types of mutations in total. In the files, you can find their corresponding mutations, as well as their swarming phenotype (branching/hyperswarming/non-swarming when growing on agar surface).

**Dataset 1**

All mutants grew in LB at 37°C with shaking for at least 12 hours. Each strain has 3 technical replicates. Growth readings were taken every 10 minutes.

**Dataset 2**

Selected mutants grew in liquid swarming medium (as defined in Luo *et al.* [28]) at 37°C with shaking. The nutrient (casamino acid) concentration was varied. Each strain has 3-6 technical replicates. There are 3 sets of data performed on different days. Growth readings were taken every 10 minutes.

Folder: rohan

**Bacterial strains**

Tetracycline resistant bacteria were evolved from ancestral strain DH5α + miniTn5-TetA under a strong promoter. Over the course of 10 days, the cells were grown in increasing tetracycline concentration. To begin, 3 μL of LB overnight culture of the ancestral strain were inoculated into 5 replicate cultures of 3 mL of LB with 2 μg/mL of tetracycline and grown overnight. Each consecutive day, 3 μL of cells were inoculated into 3 mL of LB with increasing tetracycline concentration each day: 4, 6, 8, 10, 20, 30, 40, and 50. Strains either had no plasmid, a medium-copy-number p15A plasmid, or a high-copy-number pUC plasmid. At the end of the experiment, 5 clones were isolated from each replicate culture (for a total of 25 clones) by dilution plating and frozen as glycerol stocks using a 1:1 ratio of 50% glycerol : bacterial culture.

**Bacterial growth and measurement**

Glycerol stocks of evolved clones were streaked onto LB+Tet50+Kan50 plates and grown at 37°C overnight. The streak plates were used to inoculate overnight starter culture of LB without any tetracycline. From these starter cultures, 100 μL of cells were put into a round bottom plate.

Then using 96-well plates, 2 μL of the cultures were inoculated into 200 μL of media. In each well, the tetracycline treatment was none, 20 μg/mL, or 50 μg/mL. Cells were grown in the plate reader (Tecan) for 30 hours and OD600 measurements were taken every 10 minutes.

Folder: shangying

**Bacterial strains**

Lab *E. coli* strains TOP10, MG1655, and DH5α were used.

**Protocol**

The growth media used was terrific broth (TB). Ethanol was used at varying concentrations from 0 to 6.5%. Cells were inoculated in TB with different concentrations of ethanol. The cells were then grown for 50 hours in a plate reader with OD600 readings every 10 minutes.

#### References


[1] T. Akiba, S. Sano, T. Yanase, T. Ohta, and M. Koyama, “Optuna: A Next-generation Hyperparameter Optimization Framework,” in *Proceedings of the 25th ACM SIGKDD International Conference on Knowledge Discovery & Data Mining*, in KDD ’19. New York, NY, USA: Association for Computing Machinery, July 2019, pp. 2623–2631. doi: 10.1145/3292500.3330701.

[2] A. Paszke *et al.*, “PyTorch: an imperative style, high-performance deep learning library,” in *Proceedings of the 33rd International Conference on Neural Information Processing Systems*, Red Hook, NY, USA: Curran Associates Inc., 2019, pp. 8026–8037.

[3] F. Pedregosa *et al.*, “Scikit-learn: Machine Learning in Python,” *J. Mach. Learn. Res.*, vol. 12, no. 85, pp. 2825–2830, 2011.

[4] G. Armstrong *et al.*, “Applications and Comparison of Dimensionality Reduction Methods for Microbiome Data,” *Front. Bioinforma.*, vol. 2, Feb. 2022, doi: 10.3389/fbinf.2022.821861.

[5] F. L. Gewers *et al.*, “Principal Component Analysis: A Natural Approach to Data Exploration,” *ACM Comput Surv*, vol. 54, no. 4, p. 70:1-70:34, May 2021, doi: 10.1145/3447755.

[6] C. Zhang *et al.*, “Temporal encoding of bacterial identity and traits in growth dynamics,” *Proc. Natl. Acad. Sci.*, vol. 117, no. 33, pp. 20202–20210, Aug. 2020, doi: 10.1073/pnas.2008807117.

[7] Y. Baig, H. R. Ma, H. Xu, and L. You, “Autoencoder neural networks enable low dimensional structure analyses of microbial growth dynamics,” *Nat. Commun.*, vol. 14, no. 1, p. 7937, Dec. 2023, doi: 10.1038/s41467-023-43455-0.

[8] H. R. Ma, H. Z. Xu, K. Kim, D. J. Anderson, and L. You, “Private benefit of β-lactamase dictates selection dynamics of combination antibiotic treatment,” *Nat. Commun.*, vol. 15, no. 1, p. 8337, Sept. 2024, doi: 10.1038/s41467-024-52711-w.

[9] P. Virtanen *et al.*, “SciPy 1.0: fundamental algorithms for scientific computing in Python,” *Nat. Methods*, vol. 17, no. 3, pp. 261–272, Mar. 2020, doi: 10.1038/s41592-019-0686-2.

[10] L. A. David *et al.*, “Host lifestyle affects human microbiota on daily timescales,” *Genome Biol.*, vol. 15, no. 7, p. R89, July 2014, doi: 10.1186/gb-2014-15-7-r89.

[11] H. Fujita *et al.*, “Alternative stable states, nonlinear behavior, and predictability of microbiome dynamics,” *Microbiome*, vol. 11, no. 1, Art. no. 1, Dec. 2023, doi: 10.1186/s40168-023-01474-5.

[12] “Two-stage dynamic deregulation of metabolism improves process robustness & scalability in engineered E. coli. - ScienceDirect.” Accessed: Jan. 09, 2025. [Online]. Available: https://www.sciencedirect.com/science/article/abs/pii/S109671762100149X?via%3Dihub

[13] J. N. Hennigan, R. Menacho-Melgar, P. Sarkar, M. Golovsky, and M. D. Lynch, “Scalable, robust, high-throughput expression & purification of nanobodies enabled by 2-stage dynamic control,” *Metab. Eng.*, vol. 85, pp. 116–130, Sept. 2024, doi: 10.1016/j.ymben.2024.07.012.

[14] S. Li, Z. Ye, E. A. Moreb, R. Menacho-Melgar, M. Golovsky, and M. D. Lynch, “2-Stage microfermentations,” *Metab. Eng. Commun.*, vol. 18, p. e00233, June 2024, doi: 10.1016/j.mec.2024.e00233.

[15] S. Li *et al.*, “Dynamic control over feedback regulatory mechanisms improves NADPH flux and xylitol biosynthesis in engineered *E. coli*,” *Metab. Eng.*, vol. 64, pp. 26–40, Mar. 2021, doi: 10.1016/j.ymben.2021.01.005.

[16] R. Menacho-Melgar *et al.*, “Scalable, two-stage, autoinduction of recombinant protein expression in E. coli utilizing phosphate depletion,” *Biotechnol. Bioeng.*, vol. 117, no. 9, pp. 2715–2727, 2020, doi: 10.1002/bit.27440.

[17] E. A. Moreb, Z. Ye, J. P. Efromson, J. N. Hennigan, R. Menacho-Melgar, and M. D. Lynch, “Media Robustness and Scalability of Phosphate Regulated Promoters Useful for Two-Stage Autoinduction in E. coli,” *ACS Synth. Biol.*, vol. 9, no. 6, pp. 1483–1486, June 2020, doi: 10.1021/acssynbio.0c00182.

[18] “Enabling high-throughput biology with flexible open-source automation | Molecular Systems Biology.” Accessed: Jan. 09, 2025. [Online]. Available: https://www.embopress.org/doi/full/10.15252/msb.20209942

[19] J. Schluter *et al.*, “The gut microbiota is associated with immune cell dynamics in humans,” *Nature*, vol. 588, no. 7837, pp. 303–307, Dec. 2020, doi: 10.1038/s41586-020-2971-8.

[20] R. Maddamsetti *et al.*, “Duplicated antibiotic resistance genes reveal ongoing selection and horizontal gene transfer in bacteria,” *Nat. Commun.*, vol. 15, no. 1, p. 1449, Feb. 2024, doi: 10.1038/s41467-024-45638-9.

[21] Y. Ha *et al.*, “Data-driven learning of structure augments quantitative prediction of biological responses,” *PLOS Comput. Biol.*, vol. 20, no. 6, p. e1012185, June 2024, doi: 10.1371/journal.pcbi.1012185.

[22] M. Ahmad *et al.*, “Tradeoff between lag time and growth rate drives the plasmid acquisition cost,” *Nat. Commun.*, vol. 14, no. 1, p. 2343, Apr. 2023, doi: 10.1038/s41467-023-38022-6.

[23] S. V. Aduru *et al.*, “Sub-inhibitory antibiotic treatment selects for enhanced metabolic efficiency,” *Microbiol. Spectr.*, vol. 12, no. 2, pp. e03241-23, Jan. 2024, doi: 10.1128/spectrum.03241-23.

[24] A. Palomino *et al.*, “Metabolic genes on conjugative plasmids are highly prevalent in Escherichia coli and can protect against antibiotic treatment,” *ISME J.*, vol. 17, no. 1, pp. 151–162, Jan. 2023, doi: 10.1038/s41396-022-01329-1.

[25] H. Prensky, A. Gomez-Simmonds, A. Uhlemann, and A. J. Lopatkin, “Conjugation dynamics depend on both the plasmid acquisition cost and the fitness cost,” *Mol. Syst. Biol.*, vol. 17, no. 3, p. e9913, Mar. 2021, doi: 10.15252/msb.20209913.

[26] J. H. Bethke, H. R. Ma, R. Tsoi, L. Cheng, M. Xiao, and L. You, “Vertical and horizontal gene transfer tradeoffs direct plasmid fitness,” *Mol. Syst. Biol.*, vol. 19, no. 2, p. e11300, Feb. 2023, doi: 10.15252/msb.202211300.

[27] F. Wu *et al.*, “Modulation of microbial community dynamics by spatial partitioning,” *Nat. Chem. Biol.*, vol. 18, no. 4, pp. 394–402, Apr. 2022, doi: 10.1038/s41589-021-00961-w.

[28] N. Luo *et al.*, “The collapse of cooperation during range expansion of Pseudomonas aeruginosa,” *Nat. Microbiol.*, vol. 9, no. 5, pp. 1220–1230, May 2024, doi: 10.1038/s41564-024-01627-8.

[29] E. Şimşek *et al.*, “Keystone engineering enables collective range expansion in microbial communities,” Jan. 14, 2025, *bioRxiv*. doi: 10.1101/2025.01.11.632568.

[30] E. Gullberg, L. M. Albrecht, C. Karlsson, L. Sandegren, and D. I. Andersson, “Selection of a Multidrug Resistance Plasmid by Sublethal Levels of Antibiotics and Heavy Metals,” *mBio*, vol. 5, no. 5, p. 10.1128/mbio.01918-14, Oct. 2014, doi: 10.1128/mbio.01918-14.

[31] C. Tan, P. Marguet, and L. You, “Emergent bistability by a growth-modulating positive feedback circuit,” *Nat. Chem. Biol.*, vol. 5, no. 11, pp. 842–848, Nov. 2009, doi: 10.1038/nchembio.218.

[32] M. Dzuricky, B. A. Rogers, A. Shahid, P. S. Cremer, and A. Chilkoti, “De novo engineering of intracellular condensates using artificial disordered proteins,” *Nat. Chem.*, vol. 12, no. 9, pp. 814–825, Sept. 2020, doi: 10.1038/s41557-020-0511-7.

[33] A. J. Meyer, T. H. Segall-Shapiro, E. Glassey, J. Zhang, and C. A. Voigt, “Escherichia coli ‘Marionette’ strains with 12 highly optimized small-molecule sensors,” *Nat. Chem. Biol.*, vol. 15, no. 2, pp. 196–204, Feb. 2019, doi: 10.1038/s41589-018-0168-3.

[34] P. Marguet, Y. Tanouchi, E. Spitz, C. Smith, and L. You, “Oscillations by Minimal Bacterial Suicide Circuits Reveal Hidden Facets of Host-Circuit Physiology,” *PLOS ONE*, vol. 5, no. 7, p. e11909, July 2010, doi: 10.1371/journal.pone.0011909.

[35] Z. Dai *et al.*, “Living fabrication of functional semi-interpenetrating polymeric materials,” *Nat. Commun.*, vol. 12, no. 1, p. 3422, June 2021, doi: 10.1038/s41467-021-23812-7.
[36] “EZ Rich Defined Medium.” Accessed: Jan. 23, 2025. [Online]. Available: https://www.genome.wisc.edu/resources/protocols/ezmedium.htm
[37] “Mapping single-cell responses to population-level dynamics during antibiotic treatment | Molecular Systems Biology.” Accessed: Mar. 07, 2025. [Online]. Available: https://www.embopress.org/doi/full/10.15252/msb.202211475
[38] K. N. Daeffler *et al.*, “Engineering bacterial thiosulfate and tetrathionate sensors for detecting gut inflammation,” *Mol. Syst. Biol.*, vol. 13, no. 4, p. 923, Apr. 2017, doi: 10.15252/msb.20167416.
[39] “Enzymatic assembly of DNA molecules up to several hundred kilobases | Nature Methods.” Accessed: Jan. 23, 2025. [Online]. Available: https://www.nature.com/articles/nmeth.1318
[40] N. T. Liberati, “Welcome To PA14 Transposon Insertion Mutant Library.” Accessed: Feb. 03, 2025. [Online]. Available: https://pa14.mgh.harvard.edu/cgi-bin/pa14/home.cgi
[41] N. T. Liberati *et al.*, “An ordered, nonredundant library of Pseudomonas aeruginosa strain PA14 transposon insertion mutants,” *Proc. Natl. Acad. Sci.*, vol. 103, no. 8, pp. 2833–2838, Feb. 2006, doi: 10.1073/pnas.0511100103.
[42] J. B. Xavier, W. Kim, and K. R. Foster, “A molecular mechanism that stabilizes cooperative secretions in Pseudomonas aeruginosa,” *Mol. Microbiol.*, vol. 79, no. 1, pp. 166–179, 2011, doi: 10.1111/j.1365-2958.2010.07436.x.
[43] Y. Tanouchi, A. Pai, N. E. Buchler, and L. You, “Programming stress-induced altruistic death in engineered bacteria,” *Mol. Syst. Biol.*, vol. 8, no. 1, p. 626, Jan. 2012, doi: 10.1038/msb.2012.57.
[44] S. Huang *et al.*, “Coupling spatial segregation with synthetic circuits to control bacterial survival,” *Mol. Syst. Biol.*, vol. 12, no. 2, p. 859, Feb. 2016, doi: 10.15252/msb.20156567.
[45] T. Baba *et al.*, “Construction of Escherichia coli K-12 in-frame, single-gene knockout mutants: the Keio collection,” *Mol. Syst. Biol.*, vol. 2, no. 1, p. 2006.0008, Feb. 2006, doi: 10.1038/msb4100050.
[46] A. Xia, J. Han, Z. Jin, L. Ni, S. Yang, and F. Jin, “Dual-Color Fluorescent Timer Enables Detection of Growth-Arrested Pathogenic Bacterium,” *ACS Infect. Dis.*, vol. 4, no. 12, pp. 1666–1670, Dec. 2018, doi: 10.1021/acsinfecdis.8b00129.
